# Different modification pathways for m^1^A58 incorporation in yeast elongator and initiator tRNAs

**DOI:** 10.1101/2022.12.16.520695

**Authors:** Marcel-Joseph Yared, Yasemin Yoluç, Marjorie Catala, Carine Tisné, Stefanie Kaiser, Pierre Barraud

**Affiliations:** Expression génétique microbienne, Université Paris Cité, CNRS, Institut de biologie physico-chimique, Paris, France; Department of Chemistry, Ludwig Maximilians University, Munich, Germany; Institute of Pharmaceutical Chemistry, Goethe-University, Frankfurt, Germany

**Keywords:** transfer RNA, tRNA, RNA modification, modification circuit, m1A, initiator tRNA, Pus4, Trm2, Trm6, Trm61, NMR

## Abstract

As essential components of the cellular protein synthesis machineries, tRNAs undergo a tightly controlled biogenesis process, which include the incorporation of a large number of posttranscriptional chemical modifications. Maturation defaults resulting in lack of modifications in the tRNA core may lead to the degradation of hypomodified tRNAs by the rapid tRNA decay (RTD) and nuclear surveillance pathways. Although modifications are typically introduced in tRNAs independently of each other, several modification circuits have been identified in which one or more modifications stimulate or repress the incorporation of others. We previously identified m^1^A58 as a late modification introduced after more initial modifications, such as Ѱ55 and T54 in yeast elongator tRNA^Phe^. However, previous reports suggested that m^1^A58 is introduced early along the tRNA modification process, with m^1^A58 being introduced on initial transcripts of initiator tRNA_i_^Met^, and hence preventing its degradation by the nuclear surveillance and RTD pathways. Here, aiming to reconcile this apparent inconsistency on the temporality of m^1^A58 incorporation, we examined the m^1^A58 modification pathways in yeast elongator and initiator tRNAs. For that, we first implemented a generic approach enabling the preparation of tRNAs containing specific modifications. We then used these specifically modified tRNAs to demonstrate that the incorporation of T54 in tRNA^Phe^ is directly stimulated by Ѱ55, and that the incorporation of m^1^A58 is directly and individually stimulated by Ѱ55 and T54, thereby reporting on the molecular aspects controlling the Ѱ55 → T54 → m^1^A58 modification circuit in yeast elongator tRNAs. We also show that m^1^A58 is efficiently introduced on unmodified tRNA_i_^Met^, and does not depend on prior modifications. Finally, we show that the m^1^A58 single modification has tremendous effects on the structural properties of yeast tRNA_i_^Met^, with the tRNA elbow structure being properly assembled only when this modification is present. This rationalizes on structural grounds the degradation of hypomodified tRNA_i_^Met^ lacking m^1^A58 by the nuclear surveillance and RTD pathways.

## INTRODUCTION

Transfer RNAs (tRNAs) are essential components of the cellular protein synthesis machineries, but also serve additional functions outside translation (1-4). To achieve these wide varieties of function within cells, tRNAs undergo a tightly controlled biogenesis process leading to the formation of mature tRNAs (5-8). The biogenesis of tRNAs typically includes the removal of the 5’-leader and 3’-trailer sequences from the precursor-tRNA transcripts, the addition of the 3’-CCA amino-acid accepting sequence, and the incorporation of a large number of posttranscriptional chemical modifications. These modifications occur at specific sites in a tightly controlled manner, which ensures that the tRNA biogenesis process effectively leads to the formation of functional tRNAs (9-13). All the cellular functions of tRNAs are to various extents affected by modifications. In particular, modifications in and around the anticodon are implicated in the decoding process (9,14-17), whereas modifications found in the tRNA core are collectively implicated in the folding and stability of tRNAs (18-20), making posttranscriptional modifications central in tRNA biology. Maturation defaults resulting in lack of modifications in the tRNA core may result in alternative folding (21,22), and often reduce tRNA stability, leading to the degradation of hypomodified tRNAs by the rapid tRNA decay (RTD) pathway (23-25) and the nuclear surveillance pathway (26-28).

Although modifications are typically introduced in tRNAs independently of each other, several modification circuits have been identified in which one or more modifications stimulate or repress the incorporation of another modification (11,29,30). This obviously drives a defined sequential order in the tRNA modification process. Most of the reported examples of this ordered modification process occur in the tRNA anticodon loop region (31-35), but modification circuits in the tRNA core have also been reported (36-38).

One of such circuits in the tRNA core involves modifications from the T-loop of yeast tRNAs. Using NMR spectroscopy to monitor the maturation of tRNAs in a time-resolved fashion in yeast extract (39), we previously identified a sequential order in the introduction of T54, Ψ55 and m^1^A58 in yeast tRNA^Phe^, with Ψ55 being introduced first, then T54 and finally m^1^A58 (38). Following a reverse genetic approach, we uncovered a cross-talk between these three modifications, with the m^1^A58 modification strongly depending on the two others. In a *pus4Δ* strain, lacking Ψ55, we indeed observed a severe slow-down in the introduction of both T54 and m^1^A58. Similarly, in a *trm2Δ* strain, lacking T54, we observed a slow-down in the introduction of m^1^A58 (38). In addition, we showed using liquid-chromatography coupled with tandem mass spectrometry (LC-MS/MS) that levels of m^1^A58 and T54 are affected in the *pus4Δ* and *trm2Δ* strains, in both yeast tRNA^Phe^ and in total yeast tRNAs, in a manner compatible with the cross-talks observed with NMR spectroscopy in yeast extracts. This demonstrated that these cross-talks in the T-loop are present in tRNA^Phe^ but also in other yeast tRNAs. Overall, the slow-down in the incorporation of modifications and the corresponding decrease in the modification levels observed in absence of a specific enzyme, namely in the *pus4Δ* and *trm2Δ* strains, was interpreted as a positive effect of the corresponding modification on the introduction of the other ones. We thus concluded that two modification circuits exist in the T-loop of yeast tRNAs, the long-branch Ψ55 → T54 → m^1^A58 circuit and the direct-branch Ψ55 → m^1^A58 circuit, without being able to conclude on the direct or indirect nature of the effect of Ψ55 on m^1^A58 (38).

Overall, this report on yeast tRNA^Phe^ identified m^1^A58 as a late modification, introduced after more initial modifications such as Ψ55, T54 and m^7^G46 (38). However, previous reports suggested that m^1^A58 is introduced early along the tRNA modification process in yeast, with m^1^A58 being introduced on initial pre-tRNA transcripts (6). Yeast initiator pre-tRNA_i_^Met^ lacking m^1^A58, but containing the 5’-leader and part of the 3’-trailer sequences, is indeed targeted by the nuclear surveillance and RTD pathways (26,27,40,41). In yeast tRNA_i_^Met^, the m^1^A58 modification is part of an unusual tRNA elbow structure involving non-canonical nucleotides A20, A54 and A60. This unusual substructure is assembled via an intricate network of interactions between the D- and T-loops and is likely conserved in eukaryotic initiator tRNAs (42). Altogether, these reports led to the model that m^1^A58 is introduced on pre-tRNA_i_^Met^ initial transcripts, which stabilizes the tRNA_i_^Met^ unique substructure, thereby preventing its degradation by the nuclear surveillance and RTD pathways. In addition, degradation of tRNA_i_^Met^ lacking m^1^A58 by the RTD pathway was recently shown to be conserved in the phylogenetically distant yeast species *S. pombe* and *S. cerevisiae* (41), suggesting that throughout eukaryotes the m^1^A58 modification is crucial to tRNA_i_^Met^ biology. Here, aiming to reconcile the apparent inconsistency regarding the incorporation of m^1^A58 in yeast tRNAs, namely as a late modification in elongator tRNA^Phe^ and as an early modification in initiator tRNA_i_^Met^, we decided to examine the m^1^A58 modification pathways in yeast elongator and initiator tRNAs. On the elongator tRNA^Phe^, we aimed at characterizing the molecular details related to the modification circuits present in the T-loop and involving Ψ55, T54 and m^1^A58, in order, in particular, to untangle direct from indirect effects. On the initiator tRNA_i_^Met^, we sought to investigate the introduction of m^1^A58 and its dependence on other modifications. In addition, we aimed at investigating the impact of the m^1^A58 modification on the structural properties of the tRNA_i_^Met^ elbow region. Understanding the maturation process of initiator tRNA_i_^Met^, and in particular the m^1^A58 incorporation, which has consequences on its stability and quality control is indeed crucial considering the central role of tRNA_i_^Met^ in translation initiation and hence gene expression.

For that, we first implemented a generic approach enabling the preparation of tRNAs containing specific modifications. We then used these specifically modified tRNAs to demonstrate that the incorporation of T54 in tRNA^Phe^ is directly stimulated by Ѱ55, and that the incorporation of m^1^A58 in tRNA^Phe^ is directly and individually stimulated by Ѱ55 and T54, with a remarkable cumulative effect when they are present together, thereby reporting in detail the molecular mechanisms controlling the Ψ55 → T54 → m^1^A58 modification circuit in yeast elongator tRNAs. We also show that m^1^A58 is efficiently introduced on unmodified tRNA_i_^Met^, and does not strictly need any prior modification, although m^5^C48,49 have a slight stimulatory effect on m^1^A58 incorporation. Finally, we show that the m^1^A58 single modification has tremendous effects on the structural properties of yeast tRNA_i_^Met^, with the tRNA elbow structure being properly assembled only when this modification is present. This provides a structural basis to the degradation of hypomodified tRNA_i_^Met^ lacking m^1^A58 by the nuclear surveillance and RTD pathways.

## RESULTS

### A generic approach to prepare tRNAs with specific modifications

In order to evaluate the effect of pre-existing modifications on the introduction of further modifications, tRNA samples with a single modification or a specific set of modifications need to be produced at first. For this purpose, we have implemented a generic method for preparing highly pure tRNAs with the desired modifications (Figure 1). Our approach is divided into four successive steps, namely (1) tRNA *in vitro* transcription and purification, (2) modification enzyme recombinant expression and purification, (3) *in vitro* tRNA modification reaction and modified tRNA purification, and (4) tRNA sample quality control with NMR spectroscopy (Figure 1). For introducing several modifications on a tRNA transcript, steps 3 to 5 can be reiterated on a tRNA sample already carrying modification(s).

**Figure 1:**
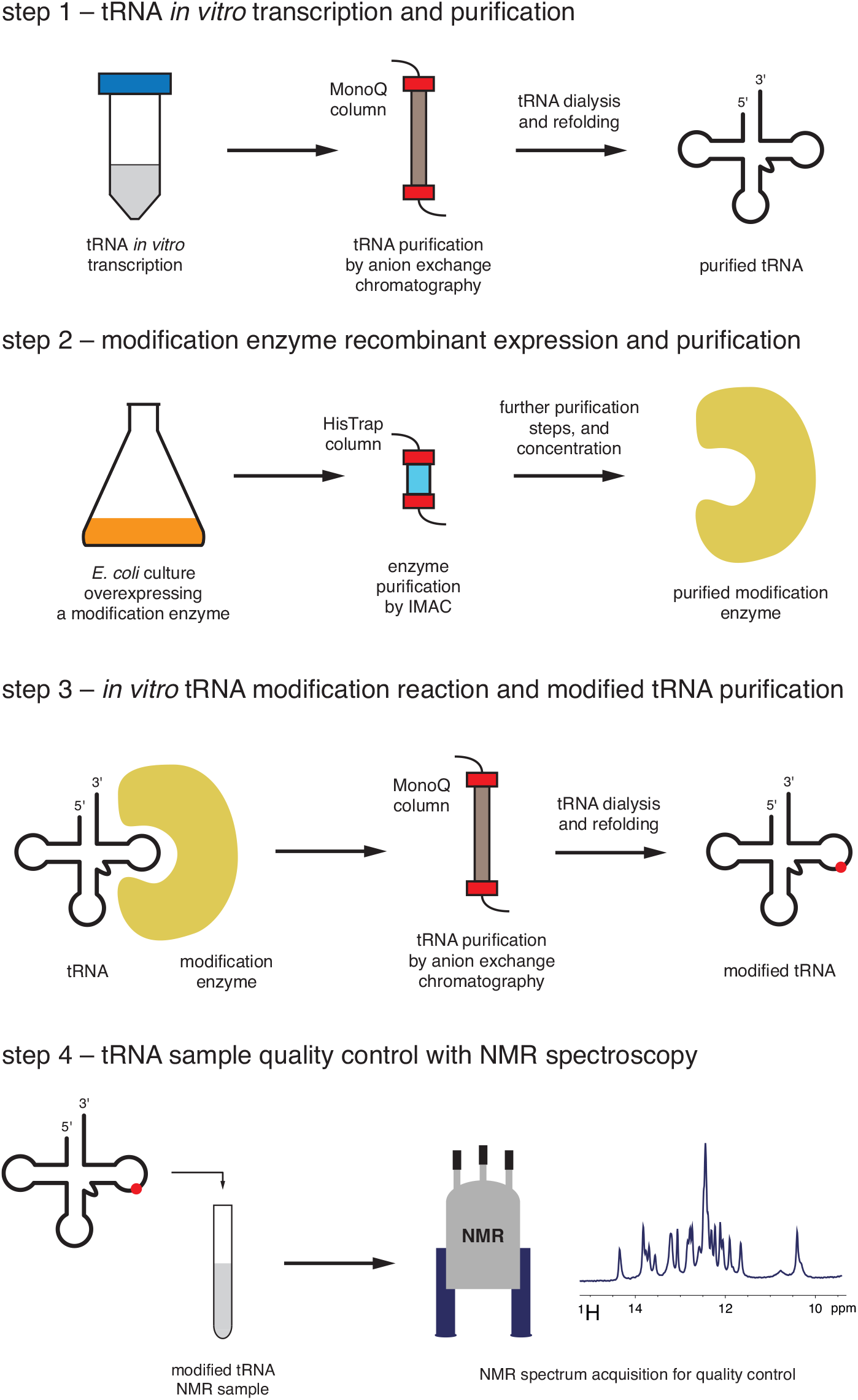
A generic approach to prepare tRNAs with specific modifications. (step 1) The unmodified tRNA is transcribed *in vitro* and purified by anion exchange chromatography. (step 2) The desired modification enzyme is overexpressed in *E. coli* and purified by immobilized metal affinity chromatography (IMAC) and further purification steps if needed. (step 3) The unmodified tRNA is modified *in vitro* with the purified modification enzyme in presence of cofactors and subsequently purified by anion exchange chromatography. (step 4) A quality control step is performed by 1D ^1^H NMR in order to establish that the desired modifications were fully incorporated in the tRNA population.

For the present study on yeast tRNA^Phe^ and tRNA_i_^Met^, in addition to the unmodified tRNA^Phe^ and tRNA_i_^Met^, we applied our methodology to produce: tRNA^Phe^ samples carrying single modifications, i.e. Ѱ55-tRNA^Phe^ and T54-tRNA^Phe^, or double modifications, i.e. Ѱ55-T54-tRNA^Phe^, and tRNA_i_^Met^ samples carrying m^5^C48,49 and m^1^A58 modifications, i.e. m^5^C48,49-tRNA_i_^Met^ and m^1^A58-tRNA_i_^Met^. For this purpose, we first transcribed and purified, using anion exchange chromatography, the yeast unmodified tRNA^Phe^ and tRNA_i_^Met^ (Figure 1, step 1). We then overexpressed and purified the yeast enzymes Pus4 that introduces Ѱ55, Trm2 that adds T54 (or m^5^U54), Trm6/Trm61 that adds m^1^A58, and Trm4 that introduces m^5^C48 and m^5^C49 (see Materials and Methods and Supplementary Figure S1 – Figure 1, step 2). Next, preliminary activity tests with these different enzymes allowed us to estimate the enzyme to tRNA ratios and the incubation times needed to introduce the desired modifications quantitatively. We thus incubated the unmodified tRNAs with the appropriate enzymes and cofactors for the required duration, and then purified the *in vitro* modified tRNAs using anion exchange chromatography (Figure 1, step 3). Finally, we verified that the desired modifications were introduced quantitatively by performing a quality control of our samples with NMR spectroscopy (Figure 1, step 4). The obtained samples are then ready to be used in downstream assays, such as activity assays of different tRNA modification enzymes.

### The introduction of T54 by Trm2 to the yeast tRNA^Phe^ is stimulated by Ѱ55

In our previous work on yeast tRNA^Phe^, we showed for a strain lacking the Ѱ55 modification, i.e. *pus4Δ* strain, that the introduction of T54 is slowed down in the *pus4Δ* yeast extract and that the amount of T54 in tRNA^Phe^ as well as in the total tRNA population is drastically reduced in the *pus4Δ* strain (38). This pointed towards a positive effect of the Ѱ55 modification in the introduction of T54 by Trm2. However, we could not exclude that the observed behaviour is due to a negative effect on the introduction of T54 caused by a certain modification or set of modifications that only become apparent in absence of Ѱ55. We could also not exclude that gene expression regulation may affect Trm2 expression in the *pus4Δ* strain. Here, in order to unambiguously determine whether the introduction of T54 on the yeast tRNA^Phe^ by Trm2 is directly dependent on the presence of the Ѱ55 modification, we conducted activity assays with Trm2 on unmodified tRNA^Phe^ and Ѱ55-tRNA^Phe^ produced as described above. Trm2 was incubated with each of the tRNAs in the presence of the methyl-donor cofactor SAM carrying a radioactive methyl group (S-adenosyl-L-methionine [methyl-^3^H]), and aliquots were taken at different time points to derive the initial velocities (Vi) of the methylation reactions (Figure 2a, Table 1, and Supplementary Figure S3). With these activity assays, we noticed that the methylation reaction catalysed by Trm2 is about 6 times faster on the Ѱ55-tRNA^Phe^ when compared to the unmodified tRNA^Phe^ (Table 2). This shows that the catalytic efficiency of Trm2 introducing T54 to tRNA^Phe^ directly depends on the prior presence of Ѱ55 and definitely establishes the direct positive link between Ѱ55 and the introduction of T54 by Trm2.

**Table 1.** Initial Velocities (Vi) of Trm2 and Trm6/Trm61 acting on yeast tRNAs presenting different modification profiles. Initial velocities were determined by linear regression and normalized to an equivalent of 50 nM of enzyme. The reported errors correspond to the standard error (SE) of the slope determination (see material and methods).

| Enzyme | Trm2 |  | Trm6/Trm61 |  |  |  |  |  |
| --- | --- | --- | --- | --- | --- | --- | --- | --- |
| Yeast tRNAs | tRNA<sup>Phe</sup> | Ψ55-tRNA<sup>Phe</sup> | tRNA<sup>Phe</sup> | T54-tRNA<sup>Phe</sup> | Ψ55-tRNA<sup>Phe</sup> | Ψ55-T54-tRNA<sup>Phe</sup> | tRNA<sub>i</sub><sup>Met</sup> | m<sup>5</sup>C-tRNA<sub>i</sub><sup>Met</sup> |
| Vi (nM/min) | 3.8 ± 0.1 | 22.9 ± 0.5 | 0.93 ± 0.05 | 3.1 ± 0.1 | 6.6 ± 0.3 | 13.7 ± 0.3 | 10.7 ± 0.3 | 15.2 ± 0.4 |

**Figure 2:**
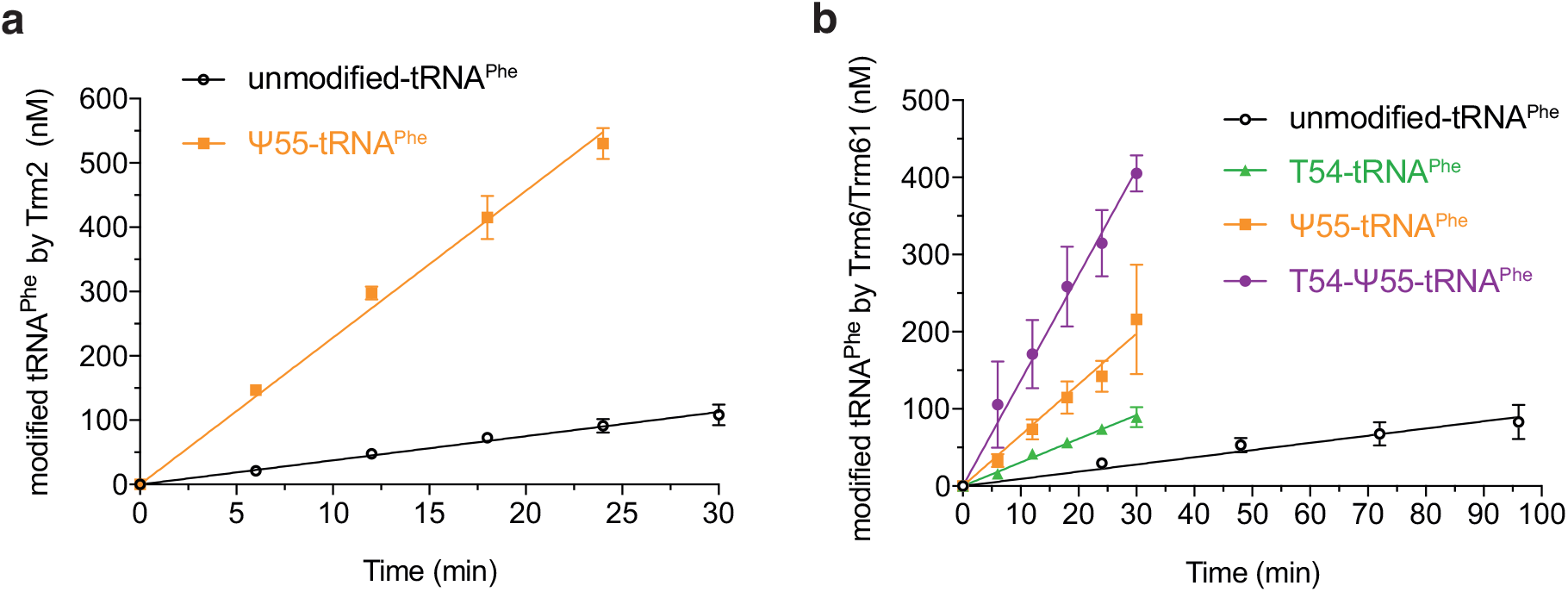
Influence of pre-existing modifications on Trm2 and Trm6/Trm61 activities on tRNA^Phe^. (**a**) Time course of the introduction of T54 in tRNA^Phe^ depending on the prior presence (orange) or absence (black) of the Ψ55 modification. (**b**) Time course of the introduction of m^1^A58 in tRNA^Phe^ depending on pre-existing modifications: unmodified tRNA^Phe^ (black), single modified T54-tRNA^Phe^ (green) and Ψ55-tRNA^Phe^ (orange), and double modified T54-Ψ55-tRNA^Phe^ (purple). Modified tRNA quantities were measured for 4 or 5 time points in at least three independent experiments (N=3 or 4), and initial velocities (Vi) were determined by linear regression (see Tables 1 and 2).

**Table 2.** Ratios of initial velocities (Vi) showing enzyme efficiency depending on the presence of pre-existing modifications on the yeast tRNA^Phe^ and tRNA_i_^Met^. The reported errors on the ratios were calculated by taking into account the propagation of uncertainties.

| Enzyme | Trm2 | Trm6/Trm61 |  |  |  |  |  |  |
| --- | --- | --- | --- | --- | --- | --- | --- | --- |
| Yeast tRNAs | Ψ55-tRNA<sup>Phe</sup> / tRNA<sup>Phe</sup> | T54-tRNA<sup>Phe</sup> / tRNA<sup>Phe</sup> | Ψ55-tRNA<sup>Phe</sup> / tRNA<sup>Phe</sup> | Ψ55-T54-tRNA<sup>Phe</sup> / tRNA<sup>Phe</sup> | m<sup>5</sup>C-tRNA<sub>i</sub><sup>Met</sup> / tRNA<sub>i</sub><sup>Met</sup> | tRNA<sub>i</sub><sup>Met</sup> / tRNA<sup>Phe</sup> | tRNA<sub>i</sub><sup>Met</sup> / Ψ55-T54-tRNA<sup>Phe</sup> | m<sup>5</sup>C-tRNA<sub>i</sub><sup>Met</sup> / Ψ55-T54-tRNA<sup>Phe</sup> |
| Vi ratio | 6.1 ± 0.2 | 3.3 ± 0.2 | 7.1 ± 0.5 | 15 ± 0.8 | 1.4 ± 0.1 | 11.5 ± 0.7 | 0.8 ± 0.03 | 1.1 ± 0.04 |

### The introduction of m^1^A58 by Trm6/Trm61 to the yeast tRNA^Phe^ is stimulated by Ѱ55 and T54

Likewise, our previous work suggested a positive effect of the Ѱ55 and T54 modifications in the introduction of m^1^A58 by the Trm6/Trm61 complex (38). However, as explained above for Trm2, we could not exclude that the observed behaviours were due to alternative effects. In addition, considering the above-mentioned effect of Ѱ55 on T54, it was not possible to distinguish whether a direct effect of Ѱ55 on m^1^A58 exists or only an indirect effect via T54. Here, in order to unambiguously determine whether the introduction of m^1^A58 on the yeast tRNA^Phe^ is directly dependent on the presence of Ѱ55 and T54, we conducted activity assays with the Trm6/Trm61 complex on unmodified tRNA^Phe^, Ѱ55-tRNA^Phe^, T54-tRNA^Phe^ and Ѱ55-T54-tRNA^Phe^ produced by following our generic approach for the production of specifically modified tRNAs. The Trm6/Trm61 complex was incubated with each of the tRNAs in the presence of a radioactive [methyl-^3^H]-SAM cofactor, and aliquots were taken at different time points to derive the initial velocities of the reactions (Figure 2b, Table 1, and Supplementary Figure S4a-d). With these activity assays, we observed that the introduction of m^1^A58 by Trm6/Trm61 is 3.3 times more efficient when the T54 modification is present compared to the unmodified tRNA^Phe^, 7.1 times more efficient in the presence of Ѱ55, and 15 times more efficient if both T54 and Ѱ55 are present in the yeast tRNA^Phe^ (Table 2). This demonstrates that T54 and Ѱ55 have individually a positive effect on the introduction of m^1^A58, as well as a cumulative positive effect if they are both simultaneously present. Therefore, the catalytic activity of Trm6/Trm61 directly depends on the presence of both the T54 and Ѱ55 modifications. Additionally, our measurements indicate that Ѱ55 stimulates the introduction of m^1^A58 about two times more efficiently than T54.

### An apparent paradox related to the essentiality of the genes encoding for the m^1^A58 modification enzyme

The results presented above showed that efficient introduction of m^1^A58 in tRNA^Phe^, strongly depends on the prior presence of the Ѱ55 and T54 modifications in the T-loop. In addition, since the observed levels of modifications in the *pus4Δ* and *trm2Δ* strains are affected in a similar manner in total yeast tRNAs and in tRNA^Phe^, the stimulation of m^1^A58 introduction by Ѱ55 and T54 is certainly a common feature of several yeast tRNAs (38). The m^1^A58 modification is introduced by the enzymatic complex Trm6/Trm61, encoded by *trm6/trm61* genes, which are among the few genes coding for tRNA modification enzymes that are essential in yeast (6,43,44).

At first sight, it might seem paradoxical that the efficiency of an enzyme encoded by two essential genes is highly dependent on the prior presence of modifications introduced by two enzymes encoded by non-essential genes. The origin of the essentiality of the m^1^A58 modification has been extensively studied and was shown to be required for the maturation and accumulation of the initiator yeast tRNA_i_^Met^ (43). Hypomodified initiator tRNA_i_^Met^ lacking m^1^A58 are targeted by the nuclear surveillance pathway and decayed by the TRAMP complex and the nuclear exosome (26,27) as well as by the RTD pathway (41). Importantly, yeast tRNA_i_^Met^ has an unusual T-loop sequence, and contains unmodified A54 and U55, which are involved in a particular tRNA elbow structure (42). We therefore anticipated that the initiator tRNA_i_^Met^ would have its own pathway of modification in the T-arm, in which the m^1^A58 modification would not depend on pre-existing modifications, and that the levels of m^1^A58 in tRNA_i_^Met^ would be unchanged in the *pus4Δ* and *trm2Δ* strains.

To demonstrate our hypothesis, we measured, using liquid-chromatography coupled with tandem mass spectrometry (LC-MS/MS), the levels of m^1^A in tRNA_i_^Met^ from *pus4Δ* and *trm2Δ* strains and compared them with the levels of m^1^A in tRNA_i_^Met^ from wild-type yeast cultured under the same experimental conditions. As expected, we do not observe any significant variation in the amount of m^1^A in the *pus4Δ* and *trm2Δ* strains when compared with the wild-type level (Figure 3a). This shows that depletion of *pus4* and *trm2* genes does not alter the activity of Trm6/Trm61 on tRNA_i_^Met^, arguing for a maintained level of expression in these strains. Therefore, in tRNA substrates carrying both Ѱ55 and T54, such as tRNA^Phe^ (38), the activity of Trm6/Trm61 is most probably primarily affected by the absence of Ѱ55 and T54, and not by the absence of the genes *pus4* and *trm2*, or gene products Pus4 and Trm2.

**Figure 3:**
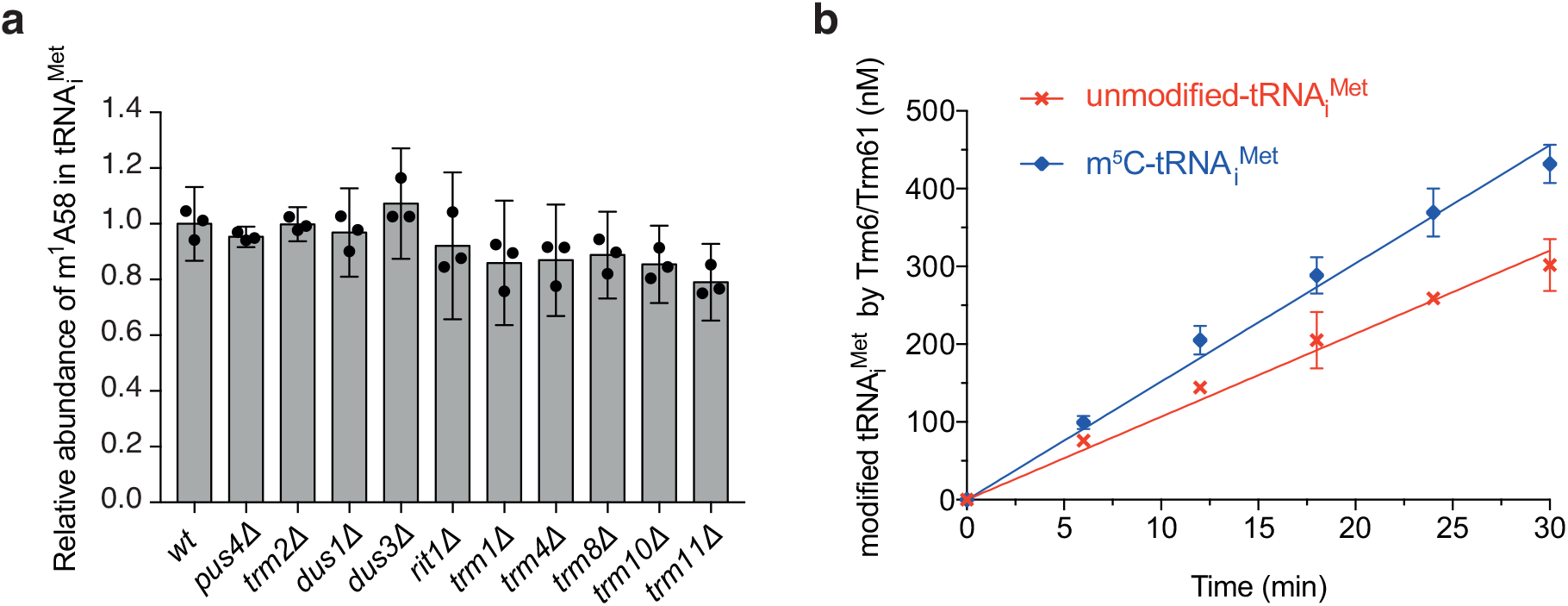
Influence of pre-existing modifications on m^1^A58 abundance and on Trm6/Trm61 activity on tRNA_i_^Met^. (**a**) Quantitative analysis of nucleoside modifications in yeast tRNA_i_^Met^ with LC–MS/MS. Histograms showing the relative abundance of m^1^A58 modification in purified yeast tRNA_i_^Met^ prepared from modification-enzyme-depleted strains using the wild-type levels as reference. Black dots represent individual measurements, data heights represent the mean of the biological replicates. Error bars correspond to the confidence interval at 95% (CI 95%). Modifications were quantified in three independent biological replicates (N=3). (**b**) Time course of the introduction of m^1^A58 in tRNA_i_^Met^ depending on the prior presence (blue) or absence (red) of the m^5^C48,49 modifications. Modified tRNA quantities were measured for 5 time points in three independent experiments (N=3), and initial velocities (Vi) were determined by linear regression (see Tables 1 and 2).

### The m^1^A58 modification is efficiently introduced on an unmodified tRNA_i_^Met^

To investigate whether any other modification present in tRNA_i_^Met^ could drive the m^1^A58 introduction, we measured similarly with LC-MS/MS the levels of m^1^A in tRNA_i_^Met^ from *dus1Δ, dus3Δ, rit1Δ, trm1Δ, trm4Δ, trm8Δ, trm10Δ* and *trm11Δ* strains, involved in the introduction of modifications D16, D47, Ar(p)64, m^2^_2_G26, m^5^C48,49, m^7^G46, m^1^G9 and m^2^G10, respectively. We observe no dramatic changes in the amount of m^1^A58 in any of these depleted strains compared with the wild-type level (Figure 3a). This suggests that m^1^A58 is efficiently introduced on yeast tRNA_i_^Met^ without the unconditional need of any prior modification. This point is perfectly in line with the m^1^A58 modification being essential for tRNA_i_^Met^ stability.

To evaluate the efficiency of m^1^A58 modification on an unmodified tRNA_i_^Met^, we conducted activity assays with Trm6/Trm61 on unmodified tRNA_i_^Met^ produced by *in vitro* transcription. The Trm6/Trm61 complex was incubated with unmodified tRNA_i_^Met^ in the presence of radioactive [methyl-^3^H]-SAM cofactor to derive the initial velocity of the reaction (Figure 3b, Table 1, and Supplementary Figure S4e). With these activity assay, we observed that the introduction of m^1^A58 by Trm6/Trm61 is 11.5 times more efficient on the unmodified tRNA_i_^Met^ than on the unmodified tRNA^Phe^ (Table 2). This corresponds to an efficiency of about 0.8 times the one measured on the doubly-modified Ѱ55-T54-tRNA^Phe^ (Table 2). Our data therefore establish that, on the contrary to its introduction on unmodified tRNA^Phe^, m^1^A58 is efficiently introduced on unmodified tRNA_i_^Met^, with an efficiency that is comparable to the one observed for an optimally modified tRNA^Phe^ bearing both Ѱ55 and T54.

### The introduction of m^1^A58 by Trm6/Trm61 to the yeast tRNA_i_^Met^ is slightly stimulated by m^5^C48,49

Even though the amount of m^1^A58 did not markedly decrease in any of the deleted strains compared with the wild-type level, some modifications may have small positive or negative effects on the kinetics of m^1^A58 introduction. Such small effects on the kinetics of introduction may not actually cause any changes in the steady-state levels of modifications in tRNAs purified from cultured yeast in the exponential phase (38). Since yeast *trm4Δ* mutant has been implicated in the RTD pathway in several combinations with other mutations (23-25), and since hypomodified tRNA_i_^Met^ are targeted by the nuclear surveillance pathway and the RTD pathway (see above), we were wondering whether the modifications introduced by Trm4 could have an impact on the introduction of m^1^A58 by Trm6/Trm61, thereby affecting tRNA_i_^Met^ stability. In yeast initiator tRNA_i_^Met^, Trm4 introduces m^5^Cs at two positions, namely m^5^C48 and m^5^C49. Another aspect that prompted us to examine the link between m^5^Cs and m^1^A58 was that m^5^C48 is very close to m^1^A58 in the tRNA_i_^Met^ structure (less than 10 Å), and that m^5^C48 and m^1^A58 are implicated in the particular tRNA elbow structure of tRNA_i_^Met^ involving the non-canonical nucleotides A20, A54 and A60 (42). More precisely, m^5^C48 is involved in an intricate network of interactions with G15, A20 and A59, with A59 and A20 forming a relay with another network involving A60 and m^1^A58 (42).

For these reasons, we decided to assess whether m^5^Cs have any effect on the introduction of m^1^A58 in tRNA_i_^Met^. We therefore conducted activity assays with Trm6/Trm61 on m^5^C48,49-tRNA_i_^Met^ produced by following our generic approach for the production of specifically modified tRNAs (Figure 1). We observe that Trm6/Trm61 is about 40% more efficient in the presence of m^5^C48,49 as compared with the unmodified tRNA_i_^Met^ (Figure 3b, Table 1, and Supplementary Figure 3f). This corresponds to an efficiency of about 1.1 times the one measured on the doubly-modified Ѱ55-T54-tRNA^Phe^ (Table 2). Thus, even though the m^5^C48,49 modifications are not strictly required for m^1^A58 introduction by Trm6/Trm61, their presence enhances the efficiency of m^1^A58 introduction in tRNA_i_^Met^ *in vitro*.

### Effect of m^5^C48,49 on the structural properties of yeast tRNA_i_^Met^

In order to better characterize the origin of m^5^Cs positive effects on m^1^A58 introduction, we undertook a structural analysis of yeast tRNA_i_^Met^ with NMR spectroscopy. To examine the consequences of m^5^C modifications on tRNA_i_^Met^, we produced both unmodified and m^5^C48,49-containing tRNA samples that are ^15^N-labelled on their imino groups, thereby allowing for the measurements of 2D ^1^H-^15^N NMR spectra, which correspond to their NMR-fingerprint and reflect their folding homogeneity and structural integrity. The comparison of the two ^1^H-^15^N BEST-TROSY experiments revealed marked differences (Figure 4a,b). Additional signals appear on the spectrum of the m^5^C48,49-tRNA, and a clear decrease in signal line broadening is observed, compared to the unmodified tRNA_i_^Met^. In addition, a pronounced signal heterogeneity is present in the unmodified tRNA_i_^Met^, with weak and strong signals coexisting, while the m^5^C48,49-tRNA_i_^Met^ spectrum shows a more homogeneous signal profile, with overall stronger signals than in the unmodified tRNA_i_^Met^ spectrum (Figure 4a,b). Since NMR signals of RNA imino groups are only observed on condition that the imino protons are protected from exchange with the solvent by hydrogen bonding in Watson-Crick or other type of base pairing, the signal heterogeneity present in the unmodified tRNA_i_^Met^ overall reflects less stable base pairs, as well as a dynamic and less homogeneous folding of this tRNA, compared with the m^5^C48,49-tRNA_i_^Met^, or the unmodified yeast tRNA^Phe^ as previously reported (45). The comparison of the NMR-fingerprints of the unmodified tRNA_i_^Met^ and the m^5^C48,49-tRNA_i_^Met^ therefore reveal that the introduction of m^5^Cs by Trm4 induces a certain stabilization in the folding of tRNA_i_^Met^, which could make it a better substrate for Trm6/Trm61 and explain the increased efficiency of m^1^A58 incorporation (Figure 3b, Tables 1 and 2).

**Figure 4:**
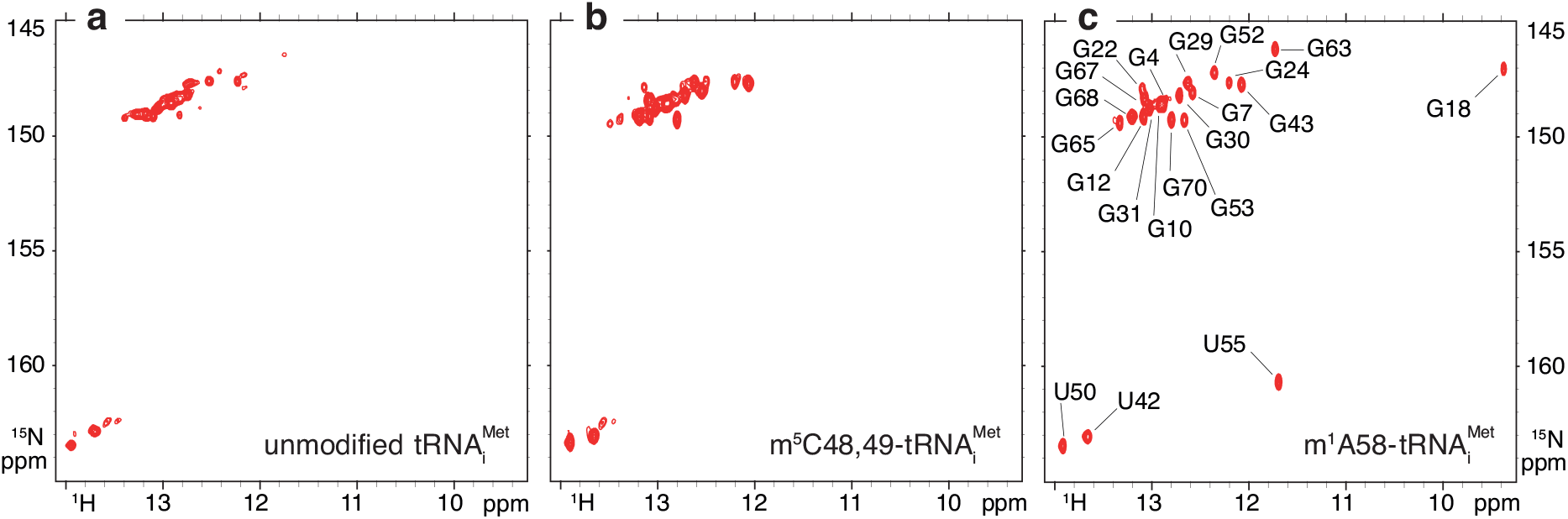
Effect of m^5^C48,49 and m^1^A58 on the structural properties of yeast tRNA_i_^Met^. Imino (^1^H,^15^N) correlation spectra of ^15^N-labelled tRNA_i_^Met^ with different modification status measured at 38°C. (**a**) unmodified tRNA_i_^Met^, **(b)** m^5^C48,49-tRNA_i_^Met^, and **(c)** m^1^A58-tRNA_i_^Met^. The assignment of the imino resonances of the m^1^A58-tRNA_i_^Met^ was obtained following standard methods.

### Effect of m^1^A58 on the structural properties of yeast tRNA_i_^Met^

Since m^1^A58 is involved in the particular tRNA elbow structure of tRNA_i_^Met^ (see reference (42) and text above), and since the m^1^A58 modification is essential for tRNA_i_^Met^ stability and prevents its degradation by the nuclear surveillance and RTD pathways (26,27,41), we were wondering what might be the effect of this single modification on the structural properties of tRNA_i_^Met^. We thus produced a ^15^N-labelled m^1^A58-containing tRNA sample following our generic approach, and measured its NMR-fingerprint (Figure 4c). The comparison of the three 1H-^15^N BEST-TROSY spectra (Figure 4a-c), and especially the comparison of the spectra of the unmodified tRNA_i_^Met^ and of the m^1^A58-tRNA_i_^Met^, revealed considerable changes in the structural properties of tRNA_i_^Met^ upon m^1^A58 modification. The pronounced signal heterogeneity present in the unmodified tRNA_i_^Met^ (Figure 4a) is completely absent in the m^1^A58-tRNA_i_^Met^ (Figure 4c), the NMR spectra of which display the characteristics of a stable and homogeneously folded tRNA. Thus, a single modification has tremendous effects on the structural properties of yeast tRNA_i_^Met^, which could likely explain why hypomodified tRNA_i_^Met^ lacking m^1^A58 are targeted by the nuclear surveillance and RTD pathways.

To get a deeper understanding of the structural changes arising upon m^1^A58 introduction, we performed the assignment of the imino resonances of the m^1^A58-tRNA_i_^Met^ following standard methods (Figure 4c), as previously described for other tRNAs (45). With this assignment at hand, we noticed that the imino signals of G18 and U55 are only visible in the spectrum of the m^1^A58-tRNA_i_^Met^ (Figure 4a-c). These nucleotides, and their respective imino groups, are engaged in universally conserved tertiary interactions at the level of the elbow region of tRNAs, with the imino group of U55 forming a hydrogen bond with a non-bridging oxygen of the phosphate backbone of A58, and that of G18 forming an hydrogen bond with an exocyclic carbonyl group of U55 (46). The detection of these imino groups in the NMR spectra of m^1^A58-tRNA_i_^Met^ attests that their imino protons are protected from an exchange with the solvent, thereby demonstrating that the tRNA elbow structure is well-assembled. The imino signals of G18 and U55 can be considered as a signature of a properly folded tRNA with a well-assembled elbow structure. Conversely, their absence in the NMR-fingerprint of the unmodified tRNA_i_^Met^ and the m^5^C48,49-tRNA_i_^Met^ (Figure 4a,b) indicate that the tRNA elbow structure is not properly assembled in these tRNAs.

### Time-resolved NMR monitoring of m^1^A58 introduction in tRNA_i_^Met^ in yeast extracts

The existence of a positive effect of m^5^Cs on m^1^A58 introduction in tRNA_i_^Met^, effect that we uncovered *in vitro* (Figure 3b), does not necessarily imply that this effect occurs in a cellular context. For instance, in the simple case where m^1^A58 would be introduced before m^5^Cs, there could not be any observable effect of m^5^Cs on the introduction of m^1^A58. In order to investigate whether this positive effect remains in a cellular context, we undertook to apply our recently developed methodology (38,39) to the monitoring of m^1^A58 introduction in tRNA_i_^Met^ in yeast extracts. As seen above, the imino signals of G18 and U55 constitute an NMR signature of a properly folded elbow structure, and therefore can be regarded as an indirect marker of m^1^A58 introduction in the case of tRNA_i_^Met^. We made use of this marker to monitor the introduction of m^1^A58 in tRNA_i_^Met^ in yeast extract. In order to assess whether m^5^Cs have a positive effect on m^1^A58 introduction in this context, we monitored, using time-resolved NMR, its incorporation in wild-type as well as in *trm4Δ* yeast extracts prepared following the same procedure. For that, ^15^N-labelled unmodified tRNA_i_^Met^ was introduced at a concentration of 40 μM in yeast extracts supplemented with the modification enzymes cofactors SAM and NADPH. The tRNA was incubated at 30°C directly in the NMR spectrometer, and a series of 1H-^15^N BEST-TROSY experiments were measured for each type of yeast extract, namely wild-type and *trm4Δ* (Figure 5). The observation of the imino signals of G18 and U55 along the tRNA_i_^Met^ maturation routes in wild-type and *trm4Δ* extracts revealed that the m^1^A58 modification is introduced slightly faster in wild-type extract than in *trm4Δ* extract depleted of Trm4. This shows that lack of m^5^C48,49 have a negative effect on m^1^A58 introduction by Trm6/Trm61 in yeast tRNA_i_^Met^, which is perfectly consistent with the *in vitro* kinetic assays on tRNA_i_^Met^ (Figure 3b). Overall, our data show that m^5^C modifications have a positive effect on m^1^A58 introduction in tRNA_i_^Met^ both *in vitro* and in a cellular context.

**Figure 5:**
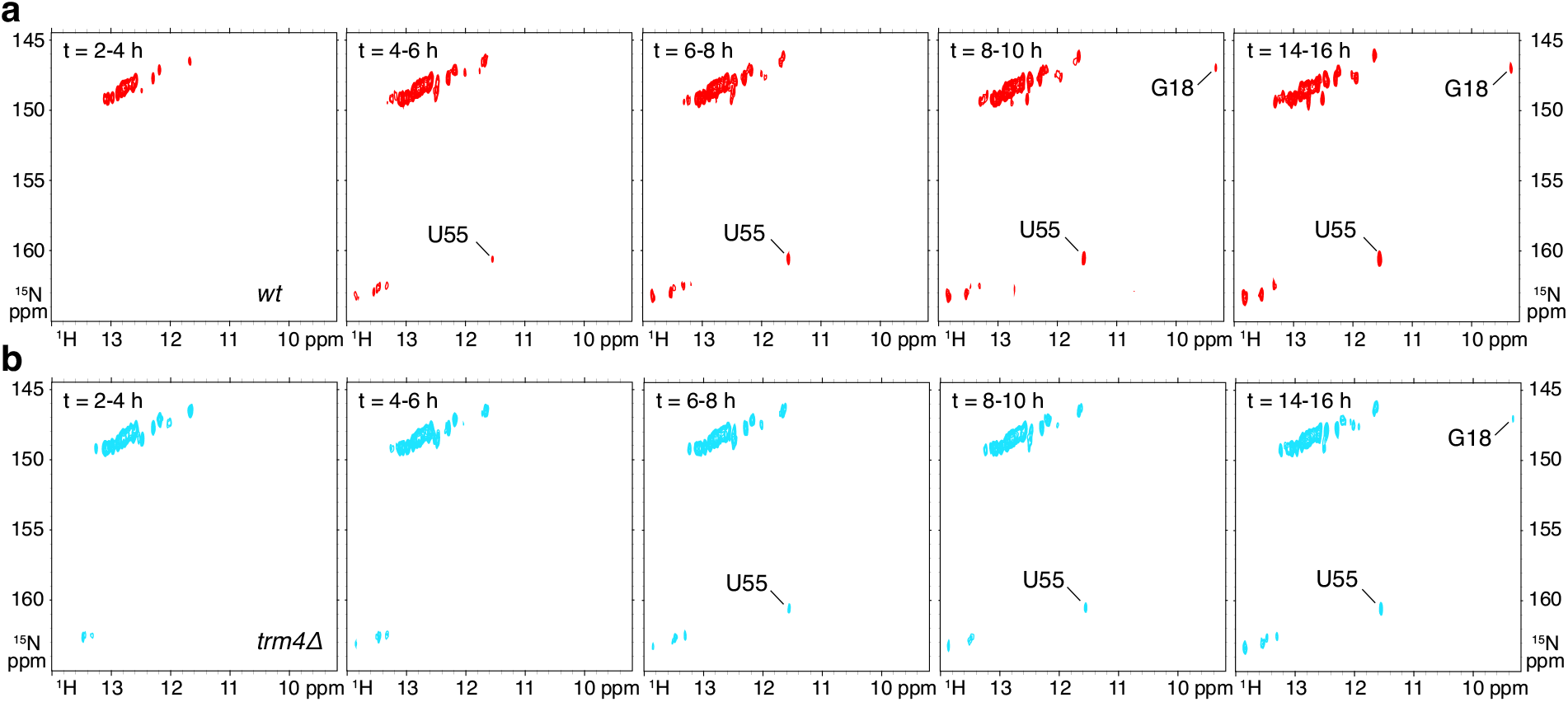
Time-resolved NMR monitoring of m^1^A58 introduction in tRNA_i_^Met^ in yeast extracts. **(a)** Imino (^1^H,^15^N) correlation spectra of a ^15^N-labelled tRNA_i_^Met^ measured in a time-resolved fashion during a continuous incubation at 30°C in yeast wild-type extract over 16 h. **(b)** Imino (^1^H,^15^N) correlation spectra of a ^15^N-labelled tRNA_i_^Met^ measured in a time-resolved fashion during a continuous incubation at 30°C in yeast *trm4Δ* extract over 16 h. Each NMR spectrum measurement spreads over a 2 h time period, as indicated.

## DISCUSSION

In this study, we implemented a generic approach for the preparation of specifically modified tRNAs in order to pursue a thorough investigation of the cross-talk between modifications Ѱ55, T54 and m^1^A58 in yeast tRNA^Phe^. We demonstrated a direct positive and cumulative effect of modifications Ѱ55 and T54 on the incorporation of m^1^A58 in this elongator tRNA. Conversely, we report that m^1^A58 is efficiently introduced on unmodified initiator tRNA_i_^Met^ without the need of any prior modification, revealing distinct pathways for m^1^A58 incorporation in yeast elongator and initiator tRNAs. Finally, we show that the m^1^A58 single modification has huge effects on the structural properties of yeast tRNA_i_^Met^, which rationalize on structural grounds the degradation of hypomodified tRNA_i_^Met^ lacking m^1^A58 by the nuclear surveillance and RTD pathways. Our study has important implications for our understanding of tRNA modification pathways and in particular for the investigation of modification circuits. These aspects are discussed below.

Genetic approaches are very effective strategies for identifying cross-talks between different genes, and genes encoding modification enzymes are no exception (29). These are however most effective when used in conjunction with biochemical approaches, allowing for a detailed characterization of the molecular aspects contributing to the observed phenotypes. With a reverse genetic approach, we previously identified an interdependence between the Ѱ55, T54 and m^1^A58 modifications in yeast tRNA^Phe^ from the observation of a slow-down in the incorporation of certain modifications in absence of other specific enzymes (38). With a biochemical approach, we now establish that the incorporation of T54 is directly stimulated by Ѱ55, and that the incorporation of m^1^A58 is directly and individually stimulated by Ѱ55 and T54, with a notable cumulative effect when they are both present, thereby reporting on direct effects of the modifications and not on other indirect effects.

A simple chemical modification, such as an isomerisation in case of Ѱ55, and a methylation in case of T54, can thus render a given tRNA a substantially better substrate for subsequent modification enzymes. The Ψ55 → T54 → m^1^A58 and Ψ55 → m^1^A58 modification circuits reported here are robust circuits with highly pronounced effects, with for instance an initial velocity of m^1^A58 incorporation that is increased by a factor 15 in presence of both Ѱ55 and T54 (Table 2). The presence of Ѱ55 alone also greatly stimulates the activity of Trm6/Trm61, with a positive effect on m^1^A58 incorporation that is about two-times larger than the positive effect of T54 (Table 2). This marked effect of Ѱ55 on the introduction of m^1^A58 lead to undetectable levels, using NMR spectroscopy, of the m^1^A58 modification along the maturation route of tRNA^Phe^ in *pus4Δ* yeast extracts (38). Similarly, m^1^A modification levels are drastically reduced in total yeast tRNAs of the *pus4Δ* strain, as reported by mass-spectrometry data (38). In addition, previous time-resolved NMR study of tRNA^Phe^ in *pus4Δ* and *trm2Δ* yeast extracts pointed towards the m^1^A58 incorporation being more affected by Ѱ55 than by T54, which is perfectly in line with the kinetic data reported here. This indicates that the time-resolved NMR approach we have developed in cellular extracts (39), is not only reliable to identify cross-talks between modifications, but also to discriminate between weak and strong dependencies.

This modification circuit in the T-arm of yeast elongator tRNAs is of interest for several reasons. First, modifications T54, Ѱ55 and m^1^A58 are among the most conserved modified nucleotides in all sequenced tRNAs, together with some dihydrouridines in the D-loop (47,48). They actually participate in maintaining the universal tRNA tertiary fold, more precisely at the level of the elbow region, assembled via conserved contacts between the T- and D-loops (46,49). Second, this modification circuit involves modified nucleotides that are found across the three domains of life, namely *Bacteria, Archaea* and *Eukarya*, and which might have been present in the prebiotic soup (50,51). Their corresponding modification enzymes probably appeared very early in evolution and were likely present in the common ancestral lineages of *Bacteria* and *Archaea* (52,53). However, all the present-day enzymes catalyzing the incorporation of T54, Ѱ55, and m^1^A58 in tRNAs, use different enzymatic strategies and/or correspond to phylogenetically unrelated enzymes between one group of organisms and another, attesting for convergent type of enzyme evolution (49). The fact that the modifications are conserved, but the corresponding enzymes are not, raises the question of whether this circuit of modifications is itself conserved between different organisms. Looking at the enzymes for which effects of prior modifications have been tested, the cross-talks as identified in yeast are likely not conserved. Effect of Ѱ55 on T54 introduction was indeed reported to be either slightly detrimental or almost neutral in full length *E. coli* tRNA^Phe^ or T-arm derived minimal stem-loop substrate (54,55). Similarly, effects of Ѱ55 and T54 on m^1^A58 introduction on a T-arm derived minimal stem-loop substrate were inconsistent with the effects reported here in yeast, with Ѱ55 being almost neutral and T54 being highly detrimental to m^1^A58 incorporation (55). Conclusions derived from measurements on minimal stem-loop substrates should however be taken with caution, since the use of a minimal substrate might alter the enzymatic activity and preference. Nevertheless, it was shown, using mass-spectrometry measurements on total yeast tRNAs, that this Ψ55 → T54 → m^1^A58 modification circuit is present in several yeast elongator tRNAs (38), and it would be of great interest to determine whether it is conserved at least among other eukaryotes.

The question remains of the molecular mechanism of such ordered modification circuit. In a circuit of modifications, the observed effect of the initial modification on the subsequent enzyme is reflected in an increased turnover rate, meaning either a better substrate binding, or a better catalytic efficiency, or a better product release, depending on the enzyme considered (56). For RNA modification enzymes, the rate-determining step of the reaction was reported to be the catalytic step (57,58), the product release (59,60), or conformational changes of both the RNA and protein, most probably to accommodate the target nucleotide into the active site (61). In a more general description, one could either consider that the initial modification directly acts as a recognition element for the subsequent modification enzyme, or that the initial modification alters the local or overall structure of the tRNA substrate, thereby presenting a structural conformation that is more efficiently modified by the subsequent enzyme. In both cases, the increased turnover rate might reflect a better substrate binding in a broad sense, which also means better substrate accommodation after a structural change via an induced-fit or conformational selection mechanism (62). The comparison of the NMR spectra of Ψ55- and T54-containing tRNA^Phe^ with that of the unmodified tRNA^Phe^, showing very limited chemical shift variations, suggests that these modifications do not induce large global rearrangements in the structure of tRNA^Phe^ (38,45). The effect of the modifications on the efficiency of the downstream enzymes is therefore most likely related to local and/or global changes in the dynamic properties of the tRNA substrate, as recently reported in the case of *E. col*i tRNAfMet, in which conformational fluctuations on the local level are increased in the modified tRNA (63). Modifications could thus help reach otherwise inaccessible structural conformations that are more suited to the subsequent modification enzyme.

Even though modification circuits are widespread and have been reported in several organisms, including *S. cerevisiae, S. pombe, E. coli, T. thermophilus*, drosophila, human and plants (31,36,37,64-70), the role of such ordered circuits of modifications remains an open question. For modification circuits in the anticodon-loop region, however, it has been recently proposed that modifications introduced first act as additional recognition elements for the subsequent enzyme, which provides the mean for adding modifications with considerable variation in the anticodon-loop region, despite the lack of variability in its local sequence and structure (30). This hypothesis is quite convincing for modifications in the anticodon-loop region, but cannot explain the actual modification circuit in the T-loop of yeast elongator tRNAs. Such a modification circuit in the tRNA core can indeed not be explained by the need for an increased variability, since the involved modifications are highly conserved in this case. Until recently, modification circuits in the tRNA core region have been only reported in the case of the hyperthermophilic bacteria *T. thermophilus* (36,37), and are most likely not implicated in sequential orders of modification incorporation, but rather in a fine tuning of modification levels in relation to an adaptation to variations in growth temperature (71). The Ψ55 → T54 → m^1^A58 modification circuit in yeast elongator tRNAs therefore constitutes the first description of an ordered circuit of modification involving modifications from the tRNA core region. Since their identification remains difficult, particularly because real-time monitoring of tRNA maturation at a single nucleotide level is technically challenging (72), we are convinced that modification circuits in the tRNA core region are certainly more widespread than currently thought.

One of the most striking features of our study concerns the spectacular changes in the structural properties of tRNA_i_^Met^ upon m^1^A58 modification. NMR-fingerprints of unmodified tRNA_i_^Met^ and m^1^A58-tRNA_i_^Met^ indeed revealed impressive structural rearrangements upon the incorporation of a single modification. Even though the NMR spectra of unmodified tRNA_i_^Met^ displays some signal heterogeneity, which attest of a certain dynamic within this tRNA that probably lead to intermediate exchange at the NMR chemical shift time-scale, the presence at the almost exact same chemical shifts of the imino groups of U42, U50, G68, G70, G12, G24 and G30 in the NMR-fingerprints of unmodified- and m^1^A58-tRNA_i_^Met^ attest of the proper secondary structure assembly of this tRNA (Figure 4). Since these imino signals could only be detected on condition that their imino protons are protected from an exchange with the solvent, all RNA helices, namely the T-, D-, anticodon-, and the acceptor-stems, are thus likely correctly assembled. However, the three-dimensional structure of the tRNA is not properly formed as demonstrated by the lack of signals attesting of a properly assembled tRNA elbow structure, namely imino groups of U55 and G18 (Figure 4). These structural rearrangements of yeast tRNA_i_^Met^ upon m^1^A58 modification are most probably at the origin of the specific degradation of hypomodified tRNA_i_^Met^ lacking m^1^A58 by the nuclear surveillance and RTD pathways (26,27,40,41), the properly folded m^1^A58-tRNA_i_^Met^ being protected from degradation. It is noteworthy that other hypomodified tRNAs lacking at least one tRNA core modification are targeted to degradation by the RTD pathway. In all reported cases, this degradation in absence of one or two modifications is tRNA specific, meaning that a specific tRNA or set of tRNAs are targeted to degradation. For instance, tRNA^Val(AAC)^ lacking m^7^G46 and m^5^C49 is rapidly degraded by the RTD pathway in *S. cerevisiae* (23); tRNA^Ser(CGA)^ and tRNA^Ser(UGA)^ lacking either Um44 and ac^4^C12 or m^2,2^G26 and m^5^C48 are also rapidly degraded by the RTD pathway in *S. cerevisiae* (25,73); tRNA^Tyr(GUA)^ and tRNA^Pro(AGG)^ lacking m^7^G46 are rapidly degraded by the RTD pathway in *S. pombe* (74). Comparing these reports with the case of tRNA_i_^Met^ lacking m^1^A58, it is tempting to speculate that the involved modifications might produce large structural effects and stabilize the tRNA tertiary structure in the particular cases of the tRNAs targeted by the RTD pathway. The same modifications would in comparison not alter much the structure of non-targeted tRNAs, a hypothesis that would need to be tested experimentally in future structural work.

Another important point revealed by the monitoring of the m^1^A58 introduction in tRNA_i_^Met^ in yeast extracts resides in the fact that our NMR-based methodology for monitoring tRNA maturation in cell extracts has the ability to report both on the introduction of chemical modifications, and on the structural changes occurring in the tRNA along the maturation pathway. This point was not fully appreciated in our former NMR study concerning yeast tRNA^Phe^, since this tRNA is to a certain extend properly folded without modifications, as evidenced by the ^1^H-^15^N NMR-fingerprint of unmodified tRNA^Phe^ that reflects a properly folded tRNA with a well-assembled elbow region (38). Changes in the NMR spectra of tRNA^Phe^ upon modification are modest (38,45), and mainly reflect the incorporation of new chemical groups, with probably also some minor structural rearrangements. The example of tRNA_i_^Met^ highlighted that NMR spectroscopy is an ideal method that can report, in a time-resolved fashion, on how the modification process affects tRNA structural properties.

In this work, we have described very different modification pathways for m^1^A58 incorporation in yeast elongator and initiator tRNAs. Unmodified elongator tRNA^Phe^ is an intrinsically poor substrate of Trm6/Trm61 for m^1^A58 incorporation, whereas unmodified tRNA_i_^Met^ is an intrinsically good substrate of the same enzyme. This raises the question of what makes a good versus a poor substrate for a modification enzyme? To look into this matter, it is important to bear in mind that modifications may not necessarily have the same beneficial effect on all tRNAs (75). For instance, a certain modification may be particularly important for a certain tRNA, which constitutes the evolutionary pressure for retaining this modification enzyme, but might be much less important, if significant at all, in other tRNAs. Such a modification would still be introduced, even though it has no actual function, due to the overlapping substrate specificity with the tRNA in which it fulfils an important function (75). Dealing with good and poor substrates therefore represents an ordinary challenge faced by modification enzymes. Indeed, tRNAs are to some extent sufficiently similar to be recognized and employed by the translation machinery, but need at the same time to be sufficiently different to be uniquely recognized by their cognate aminoacyl-tRNA synthetases. The modification enzymes therefore should handle a population of highly similar but unique tRNAs and the tRNA modification patterns can be regarded as the result of million years of coevolution of modification enzymes with the tRNA population (11,76). In this context, we believe that a potential role of modification circuits could be to allow the modification of both good and poor tRNA substrates. In the case of yeast elongator and initiator tRNAs, that must have sufficiently different structural properties to be recognized by elongation or initiation factors, respectively, the existence of the Ψ55 → T54 → m^1^A58 modification circuit enables the incorporation of m^1^A58 in certain elongator tRNAs, such as tRNA^Phe^, which are poor intrinsic substrate of Trm6/Trm61. In conclusion, modification circuits might be a solution found to deal with the problem of having at the same time poor and good tRNA substrates that all require to be eventually modified.

## METHODS

### Yeast strains

Yeast strains used in this study are listed in Supplementary Table S1. The wild-type *S. cerevisiae* BY4741 strain and the YKO collection kanMX strains carrying deletions of the genes for modification enzymes Pus4, Trm1, Trm2, Trm4, Trm8, Trm10, Trm11, Pus4, Dus1, Dus3, and Rit1, were obtained from Euroscarf and used for tRNA preparations for MS analysis. The proteinase-deficient *S. cerevisiae* strain c13-ABYS-86 and the derived strain c13-ABYS-86-*trm4Δ* were used for the preparation of yeast extracts for NMR experiments. All strain constructions were verified by PCR using appropriate oligonucleotides (listed in Supplementary Table S2).

### *E. coli* strains

*E. coli* strains used in this study are listed in Supplementary Table S1. The *E. coli* BL21(DE3) CodonPlus-RIL *yggh::kan* strain was constructed by transferring the *yggh::kan* cassette from the appropriate K-12 strain of the Keio collection (77) to a BL21(DE3) CodonPlus-RIL strain (Agilent) by phage P1 *vir*-mediated transduction (78) (Supplementary Table S1). Deletion of the *yggh* gene and its replacement by the kanamycine resistance cassette in the BL21(DE3) CodonPlus-RIL strain was checked with PCR using appropriate sets of primers (Supplementary Table S2).

### Pus4 modification enzyme cloning and purification

The gene encoding the full-length yeast Pus4 (M1 to V403 – Uniprot entry P48567) was cloned from BY4741 genomic DNA between the *EcoRI* and *NotI* sites of a modified pET28a vector (Novagen) encoding an N-terminal His6-tag cleavable with TEV protease (pET28-Pus4). The overexpression and purification procedures were adapted from a previously published procedure (79). Briefly, Pus4 was overexpressed in *E. coli* BL21(DE3) CodonPlus-RIL cells (Agilent) in LB media. The cells were grown at 37°C to OD600 ∼0.4, cooled down at 30°C and induced at OD600 ∼0.6 by adding isopropyl-β-D-thiogalactopyranoside (IPTG) to a final concentration of 0.5 mM. Cells were harvested 6 h after induction by centrifugation. Cell pellets were resuspended in lysis buffer (50 mM Tris-HCl pH 8.0, 300 mM NaCl, 10 mM 2-mercaptoethanol, 1 mM phenylmethanesulfonylfluoride (PMSF), 10% (v/v) glycerol, 1 mM ethylenediaminetetraacetic acid (EDTA)) supplemented with an EDTA-free antiprotease tablet (Roche) and lysed by sonication. Cell lysates were centrifuged for 30 min at 35,000 g. All column chromatography purifications were performed on a ÄKTA Pure purification system (Cytiva) at 4°C. The cell lysate supernatant was loaded on a Ni-NTA column and the protein of interest was eluted with an imidazole gradient. Fractions containing the protein were pooled, concentrated with an Amicon 50,000 MWCO (Millipore) and further purified with a Superdex 75 size exclusion column (Cytiva). Purified proteins were eluted in the Pus4 storage buffer (50 mM Tris-HCl pH 8.0, 150 mM NaCl, 2 mM 2-mercaptoethanol), confirmed for purity using SDS-PAGE (Supplementary Figure S1), concentrated with an Amicon 10,000 MWCO (Millipore) to ∼10 mg/mL and stored at -20°C. The protein concentration was determined by absorbance at 280 nm using a mM extinction coefficient of 30.4 mM^-1^.cm^-1^.

### Trm2 modification enzyme cloning and purification

The gene encoding the full-length yeast Trm2 (M1 to I639 – Uniprot entry P33753) was initially cloned from BY4741 genomic DNA between the *EcoRI* and *NotI* sites of a pGEX-6p-1 vector (pGEX-Trm2). However, this construct happened to be insoluble and poorly expressed in *E. coli* BL21(DE3) CodonPlus-RIL cells (Agilent). Since the N-terminal part of Trm2 contains highly hydrophobic stretches of amino-acids, and does not correspond to the catalytic domain of the protein, a second construct corresponding to the yeast Trm2 V116 to I639 residues was cloned between the *BamHI* and *XhoI* sites of a pRSFDuet-Smt3 vector leading to an N-terminal His6-SUMO-fusion of Trm2 (pSUMO-Trm2p). The naturally present *BamHI* site into the yeast *trm2* gene was first removed by a silent mutation of the codon encoding for D564 from GAT to GAC with site directed mutagenesis. The yeast Trm2 protein was overexpressed in *E. coli* BL21(DE3) CodonPlus-RIL cells in LB media. The cells were grown at 37°C to OD600 ∼0.4, cooled down at 25°C and induced at OD600 ∼0.6 by adding IPTG to a final concentration of 0.5 mM. Cells were harvested 6 h after induction by centrifugation. Cell pellets were resuspended in lysis buffer (50 mM Tris-HCl pH 8.0, 1 M NaCl, 5 mM MgCl2, 1 mM dithiothreitol (DTT), 5% (v/v) glycerol, 1 mM EDTA) supplemented with an EDTA-free antiprotease tablet (Roche) and lysed by sonication. Cell lysates were centrifuged for 30 min at 35,000 g. All column chromatography purifications were performed on a ÄKTA Pure purification system (Cytiva) at 4°C. The cell lysate supernatant was loaded on a Ni-NTA column and the protein of interest was eluted with an imidazole gradient. The fractions containing the protein were pooled, supplemented with ammonium sulfate then loaded on a HiPrep Phenyl HP 16/10 hydrophobic Column (Cytiva) and eluted with a decreasing ammonium sulfate gradient. Fractions containing the protein were pooled, concentrated with an Amicon 50,000 MWCO (Millipore) and further purified with a Superdex 200 size exclusion column (Cytiva). Purified proteins were eluted in the Trm2 storage buffer (50 mM Tris-HCl pH 8.0, 400 mM NaCl, 1 mM DTT), confirmed for purity using SDS-PAGE (Supplementary Figure S1), concentrated with an Amicon 50,000 MWCO (Millipore) to ∼5 mg/mL and stored at -20°C. The protein concentration was determined by absorbance at 280 nm using a mM extinction coefficient of 30.8 mM^-1^.cm^-1^ for the SUMO-Trm2 construct.

### Trm6/Trm61 modification enzyme cloning and purification

The genes encoding yeast Trm6/Trm61 heterodimer (Trm6: M1 to I478 – Uniprot entry P41814; Trm61: M1 to K383 – Uniprot entry P46959) were cloned from BY4741 genomic DNA between the *BamHI* and *NotI* sites for Trm6 and *NdeI* and *XhoI* sites for Trm61 of a pETDuet-1 vector (Novagen) thereby encoding an N-terminal His6-tag on Trm6 (pETDuet-Trm6/Trm61). Since initial expression and purification in *E. coli* BL21(DE3) CodonPlus-RIL cells (Agilent) lead to a Trm6/Trm61 heterodimer contaminated with an m^7^G46 modification activity (see Supplementary Figure S2), subsequent expressions of the Trm6/Trm61 heterodimer were conducted in *E. coli* BL21(DE3) CodonPlus-RIL *yggh::kan* cells lacking the *E. coli* enzyme catalyzing m^7^G46 modifications in tRNAs, namely TrmB. The overexpression and purification procedures were adapted from a previously published procedure (80). The Trm6/Trm61 protein complex was overexpressed in *E. coli* BL21(DE3) CodonPlus-RIL *yggh::kan* cells in LB media. The cells were grown at 37°C to OD600 ∼0.4, cooled down at 18°C and induced at OD600 ∼0.6 by adding IPTG to a final concentration of 0.4 mM. Cells were harvested 22 h after induction by centrifugation. Cell pellets were resuspended in lysis buffer (50 mM Tris-HCl pH 8.0, 300 mM NaCl, 1 mM PMSF, 1 mM DTT, 5% (v/v) glycerol, 1 mM EDTA) and lysed by sonication. Cell lysates were centrifuged for 30 min at 35,000 g. All column chromatography purifications were performed on a ÄKTA Pure purification system (Cytiva) at 4°C. The cell lysate supernatant was loaded on a Ni-NTA column and the protein of interest was eluted with an imidazole gradient. Fractions containing the protein were pooled, concentrated with an Amicon 50,000 MWCO (Millipore) and further purified with a Superdex 200 size exclusion column (Cytiva). Purified proteins were eluted in the Trm6/Trm61 storage buffer (50 mM Tris-HCl pH 8.0, 300 mM NaCl, 5mM DTT), confirmed for purity using SDS-PAGE (Supplementary Figure S1), concentrated with an Amicon 50,000 MWCO (Millipore) to ∼5 mg/mL and stored at -20°C. The protein concentration was determined by absorbance at 280 nm using mM extinction coefficients of 178 mM^-1^.cm^-1^ for the Trm6/Trm61 heterotetramer.

### Trm4 modification enzyme purification

The gene encoding the full-length yeast Trm4 (M1 to N684 – Uniprot entry P38205) was overexpressed from a pET28-Trm4 vector and purified following procedures adapted from a previously published protocol (81). Briefly, the yeast Trm4 protein was overexpressed in *E. coli* BL21(DE3) CodonPlus-RIL cells in LB media. The cells were grown at 37°C to OD600 ∼0.4, cooled down at 30°C and induced at OD600 ∼0.6 by adding IPTG to a final concentration of 0.5 mM, they were then left to grow overnight. Cells were harvested ∼17 h after induction by centrifugation. Cell pellets were resuspended in lysis buffer (50 mM Tris-HCl pH 8.0, 0.5 M NaCl, 1 mM dithiothreitol (DTT), 0.1% (v/v) Triton X-100, 1 mM EDTA), supplemented with an EDTA-free antiprotease tablet (Roche) and lysed by sonication. Cell lysates were centrifuged for 30 min at 35,000 g. All column chromatography purifications were performed on an ÄKTA Pure purification system (Cytiva) at 4°C. The cell lysate supernatant was loaded on a Ni-NTA column and the protein of interest was eluted in an imidazole gradient. Fractions containing the protein were pooled, concentrated with an Amicon 50,000 MWCO (Millipore) and further purified with a Superdex 200 size exclusion column (Cytiva). Purified proteins were eluted in the Trm4 storage buffer (50 mM Tris-HCl pH 8.0, 300 mM NaCl, 1 mM DTT, 5% (v/v) glycerol), confirmed for purity using SDS-PAGE (Supplementary Figure S1), concentrated with an Amicon 50,000 MWCO (Millipore) to ∼8 mg/mL and stored at -20°C. The protein concentration was determined by absorbance at 280 nm using a mM extinction coefficient of 80.9 mM^-1^.cm^-1^.

### RNA sample preparation for NMR and kinetic assays

Unmodified yeast tRNA^Phe^ and tRNA_i_^Met^ were prepared by standard *in vitro* transcription following previously published procedures, either with unlabelled NTPs or ^15^N-labelled Us and Gs (38,82). We replaced the first Watson Crick base pair A1-U72 of the tRNA_i_^Met^ with a G1-C72 base pair in order to improve *in vitro* transcription efficiency. To prepare the single modified Ψ55-tRNA^Phe^, 112 μM of refolded tRNA^Phe^ was incubated with 3.3 μM of purified Pus4 for 40 min at 30°C in an 800 μL reaction mix. To prepare T54-tRNA^Phe^, 80 μM of refolded tRNA^Phe^ was incubated with 12 μM of purified Trm2 and a ∼6-8-times excess of S-adenosyl-L-methionine (SAM) in an 800 μL reaction mix for 14 h at 30°C. To prepare the double modified Ψ55-T54-tRNA^Phe^, 80 μM of Ψ55-tRNA^Phe^ was incubated with 8 μM Trm2 and a ∼6-8-times excess of SAM in an 800 μL reaction mix for 4 h at 30°C. To prepare m^5^C48,49-tRNA_i_^Met^, 146 μM of refolded unmodified-tRNA_i_^Met^ was incubated with 23 μM of purified Trm4 for 17 h at 30°C in a 500 μL reaction mix. All reactions were performed in the following maturation buffer (MB): 100 mM NaH2PO4/K2HPO4 pH 7.0, 5 mM NH4Cl, 2 mM DTT and 0.1 mM EDTA. The tRNA reaction products were then purified by ion exchange chromatography (MonoQ, Cytiva), dialyzed extensively against 1 mM Na-phosphate pH 6.5, and refolded by heating at 95°C for 5 min and cooling down slowly at room temperature. Buffer was added to place the tRNAs in the NMR buffer (10 mM Na-phosphate pH 6.5, 10 mM MgCl2), and the samples were concentrated using Amicon 10,000 MWCO (Millipore) to ∼80 μM for further use in kinetic assays, or ∼1.4-1.5 mM for the NMR study of tRNA_i_^Met^ maturation in yeast extracts.

### Enzyme kinetic assays

To measure the m^1^A58 formation, 10 μM of unmodified tRNA^Phe^, Ψ55-tRNA^Phe^, T54-tRNA^Phe^, Ψ55-T54-tRNA^Phe^, unmodified tRNA_i_^Met^ and m^5^C48,49-tRNA_i_^Met^ were incubated each in a 300 μL reaction with 300 nM, 100 nM, 300 nM, 50 nM, 150 nM, and 150 nM of purified Trm6/Trm61, respectively, 18 μM non-radioactive SAM and 50 nM of radioactive [^3^H]-SAM (Supplementary Table S3). To measure the T54 formation, 10 μM of unmodified tRNA^Phe^ and Ψ55-tRNA^Phe^ were incubated each in a 300 μL reaction with 400 nM and 50 nM of Trm2, respectively, 18 μM non-radioactive SAM and 50 nM of radioactive [^3^H]-SAM (Supplementary Table S3). Reactions were performed in the MB buffer and were incubated at 30°C for 30 min except for the reaction with Trm6/Trm61 and the unmodified tRNA^Phe^ that was incubated for 96 min. Aliquots of 50 μL were taken of each reaction at 6, 12, 18, 24 and 30 min (for the 30 minutes reactions) and at 24, 48, 72 and 96 min (for the 96 min reaction) and the samples were quenched by adding 5% (v/v) cold trichloracetic acid (TCA). Quenched samples were filtered through Whatman glass microfibers disks pre-soaked with 5% (v/v) TCA, washed four times with 5% (v/v) TCA, and one final time with ethanol. The filter disks were dried, then 5 mL Optiphase ‘HISAFE’ 2 scintillation cocktail (PerkinElmer) were added, and the counts per minute (CPM) equivalent to the incorporated [^3^H]-methyl were determined by scintillation counting. Then CPM values were converted to concentrations of modified tRNAs using [^3^H]-SAM/CPM calibration standards. Enzymatic reactions were performed in triplicates or quadruplicates. Since Trm6/Trm61 and Trm2 activity turned out to vary greatly between different substrates, different enzyme concentrations were used to perform the kinetic assays. Therefore, we normalized the quantities of modified tRNAs to an equivalent of 50 nM of enzyme. Initial velocities (V_i_) were determined by linear regression using Prism7 (GraphPad), *i*.*e*. data were fitted to a single linear function: y= V_i_ .x while forcing the curve to pass through the origin, and standard errors (SE) on the V_i_ were determined by taking into account the data spread.

### NMR spectroscopy

All NMR spectra of yeast tRNA^Phe^ and tRNA_i_^Met^ were measured at 38°C on a Bruker AVIII-HD 700 MHz spectrometer equipped with TCI 5-mm cryoprobe with 5-mm Shigemi tubes in the NMR buffer (10 mM Na-phosphate pH 6.5, 10 mM MgCl2) supplemented with 5% (v/v) D2O. To verify that the desired modifications were incorporated quantitatively in yeast tRNA^Phe^, 1D jump-and-return-echo NMR spectra (83,84) of the different tRNAs were measured, and compared to previously characterized samples (38,45). In addition, to evaluate the effect of specific modifications on the structural properties of yeast tRNA_i_^Met^, 2D (^1^H,^15^N)-BEST-TROSY spectra of unmodified tRNA_i_^Met^, m^5^C48,49-tRNA_i_^Met^ and m^1^A58-tRNA_i_^Met^ were measured at 38°C in the NMR buffer. Imino resonances of the m^1^A58-tRNA_i_^Met^ were assigned using 2D jump-and-return-echo (^1^H,^1^H)-NOESY (83,84) and 2D (^1^H,^15^N)-BEST-TROSY (85) experiments. For monitoring the maturation of tRNA_i_^Met^ in yeast extract, wild-type and *trm4Δ* yeast extracts were prepared in the c13-ABYS-86 background, as previously described (39). NMR spectra were measured at 30°C with unmodified ^15^N-[U/G]-labelled tRNA_i_^Met^ at 40 μM in yeast extracts supplemented with NaH2PO4/K2HPO4 pH 6.5 150 mM, NH4Cl 5 mM, MgCl2 5 mM, DTT 2 mM, EDTA 0.1 mM, SAM 4 mM, ATP 4 mM, NADPH 4 mM, and D2O 5% (v/v) (86). Each 2D (^1^H,^15^N)-BEST-TROSY experiment of the series was measured with a recycling delay of 200 ms, a SW(^15^N) of 26 ppm, and 96 increments for a total experimental time of 120 min. The data were processed using TOPSPIN 3.6 (Bruker) and analysed with Sparky (http://www.cgl.ucsf.edu/home/sparky/).

### Total tRNA samples from yeast for mass spectrometry

Total tRNA from *S. cerevisiae* BY4741 wild-type or mutant strains used for mass spectrometry analysis were prepared as described previously (38). For each strain, all cultures and tRNA preparations were performed in triplicate for statistical analysis. Yeast tRNA_i_^Met^ was isolated from ∼1 μg total tRNA samples with a first step of SEC and a subsequent purification using T1 Dynabeads (Thermo Fisher Scientific, Product no. 65801D) and a DNA probe specific to tRNA_i_^Met^ ([Btn]-AAA-TCG-GTT-TCG-ATC-CGA-GGA-CAT-CAG-GGT-TAT-GA, Sigma-Aldrich, Munich, Germany) as previously reported (38,87,88).

### Digestion of tRNAs to nucleosides and quantification by mass spectrometry

Specifically purified to tRNA_i_^Met^ samples were digested to single nucleosides following previously published procedures (38) and stable isotope-labelled internal standard (SILIS, 0.1 volume of 10X solution) from yeast was added for absolute quantification (89). Quantification of the m^1^A modification in tRNA_i_^Met^ was performed with an Agilent 1290 Infinity II equipped with a DAD combined with an Agilent Technologies G6470A Triple Quad system and electrospray ionization (ESI-MS, Agilent Jetstream) following previously published procedures (38,89). Analyses of the variations compared to the wild-type strain were conducted from the determination of the confidence intervals at 95% (CI 95%) using Prism7 (GraphPad).

## AUTHORS CONTRIBUTION

PB conceived the study. PB cloned the constructs for Pus4, Trm2 and Trm6/Trm61 overexpression and constructed the *E. coli* BL21 *yggh::kan* strain. MJY purified all modification enzymes with support from MC and under the supervision of PB and CT. MJY prepared tRNA samples for NMR and kinetic assays. MJY performed and analyzed the enzymatic assays. MJY prepared total tRNA samples for MS; YY isolated and digested tRNA_i_^Met^ for MS; YY measured and analyzed MS data under the supervision of SK. MJY and PB measured and analyzed the NMR spectra. MJY and PB wrote the manuscript.

## ACKNOWLEDGEMENTS

The authors are grateful to Josette Banroques for providing the yeast YKO kanMX strains, to Grégory Boël for providing the *E. coli* K-12 *yggh::kan*, to Julien Henri for providing the pRSFDuet-Smt3 vector, to Sylvie Auxilien and Yuri Motorin for providing the pET28-Trm4 expression plasmid, to Alexandre Gato for preliminary data acquisition, and to Christel Le Bon for ensuring the best performance of the NMR infrastructure at the IBPC. The authors acknowledge access to the biomolecular NMR platform of the IBPC that is supported by the CNRS, the Labex DYNAMO (ANR-11-LABX-0011), the Equipex CACSICE (ANR-11-EQPX-0008) and the Conseil Régional d’Île-de-France (SESAME grant). This work was funded by the ANR JCJC CiMoDyMo (ANR-19-CE44-0013), the Deutsche Forschungsgemeinschaft (255344185-SPP 1784) and the Horizon 2020 program (ID-952373).

