## Supplementary Material for "Different modification pathways for m^1^A58 incorporation in yeast elongator and initiator tRNAs"

---

Content:

| | |
|---|---|
| Supplementary Tables S1 to S3 | pages 2-4 |
| Supplementary Figures S1 to S4 | pages 5-8 |
| Supplementary References | page 9 |

### SUPPLEMENTARY TABLES

#### Supplementary Table S1 – Yeast and *E. coli* strains used in this study

| Strain | Genotype | Source |
|---|---|---|
| BY4741 | *MATa his3-Δ1 leu2-Δ0 met15-Δ0 ura3-Δ0* | Euroscarf |
| BY4741-*dus1Δ* | BY4741, *YML080w::kanMX* | Euroscarf |
| BY4741-*dus3Δ* | BY4741, YLR401c*::kanMX* | Euroscarf |
| BY4741-*pus4Δ* | BY4741, *YNL292w::kanMX* | Euroscarf |
| BY4741-*trm1Δ* | BY4741, *YDR120c::kanMX* | Euroscarf |
| BY4741-*trm2Δ* | BY4741, *YKR056w::kanMX* | Euroscarf |
| BY4741-*trm4Δ* | BY4741, *YBL024w::kanMX* | Euroscarf |
| BY4741-*trm8Δ* | BY4741, *YDL201w::kanMX* | Euroscarf |
| BY4741-*trm10Δ* | BY4741, *YOL093w::kanMX* | Euroscarf |
| BY4741-*trm11Δ* | BY4741, *YOL124c::kanMX* | Euroscarf |
| BY4741-*rit1Δ* | BY4741, *YMR283c::kanMX* | Euroscarf |
| c13-ABYS-86 | *MATα ura3Δ5 leu2-3,112 his3 pra1-1 prb1-1 prc1-1 cps1-3* | (1) |
| c13-ABYS-86-*trm4Δ* | c13-ABYS-86, *YBL024w::kanMX* | (2) |
| BW25113-*ygghΔ* | *E. coli* K-12 $lacI^{+}$ $rrnB_{T14}$ $\Delta lacZ_{WJ16}$ *hsdR*514 $\Delta araBAD_{AH33}$ $\Delta rhaBAD_{LD78}$ *rph-1* Δ*(araB–D)567* Δ*(rhaD–B)568* Δ*lacZ4787*(::*rrnB-3*) *hsdR514 rph-1 yggh::kan* | (3) |
| BL21-CodonPlus (DE3)-RIL | *E. coli* B $F^{-}$ ompT hsdS($r_B^{-}m_B^{-}$) $dcm^{+}$ $Tet^{r}$ gal λ(DE3) endA Hte [argU ileY leuW $Cam^{r}$] | Agilent |
| BL21-CodonPlus (DE3)-RIL-*ygghΔ* | *E. coli* BL21-CodonPlus(DE3)-RIL *yggh::kan* | This study |

**Supplementary Table S2 – List of primers used in this study**

Pus4-M1_EcoRI_Fwd: CAG-GCG-AAT-TCA-TGA-ATG-GAA-TAT-TTG-CTA-TT
Pus4-V403_NotI_Rev: TAC-AAG-GCG-GCC-GCT-TAC-ACC-TGT-TCG-ATT-TT

Trm2-M1_EcoRI_Fwd: GAC-AAA-GAA-TTC-ATG-TAC-GAA-CAG-TTT-GAA-TTT
Trm2-I639_NotI_Rev: TAT-TTT-GCG-GCC-GCA-ATT-AGA-TTC-TCT-TCA-TTA
Trm2_BamHI-del_Fwd: GTT-ATC-TTG-GAC-CCA-CCA-CGC-AAG-GGC-TGT-GAC-GAA-TTA-TTC
Trm2_BamHI-del_Rev: TGG-TGG-GTC-CAA-GAT-AAC-GGA-AGT-GTT-TTC-ACT-TGG-AGT-ATC-AAT-AG
Trm2-V116_BamHI_Fwd: CGA-TGG-ATC-CGT-TGA-AAC-AAC-TTC-TCC-GAT-GG
Trm2-I639_XhoI_Rev: GCG-CCT-CGA-GTT-AGA-TTC-TCT-TCA-TTA-TAC-ACA-C

Trm6-M1_BamHI_Fwd: AAT-TTA-GGA-TCC-GAT-GAA-TGC-TTT-GAC-AAC
Trm6-I478_NotI_Rev: CTC-TGC-GGC-CGC-TTA-TAT-CTT-TTG-TTT-CTT-AG
Trm61-M1_NdeI_Fwd: TTT-ACA-TAT-GTC-AAC-AAA-TTG-TTT-TTC-CGG-TTA
Trm61-K383_XhoI_Rev: TAA-TCC-TCG-AGT-TAT-TTT-TCC-GTG-GAT-CGA-AGA

yggh_A: GGT-GCG-TAC-CTC-ATC-CAG-TT
yggh_B: GTA-TCG-TTC-AGG-TGC-CGT-TT
yggh_C: CTG-CTC-AGG-GCG-ATC-TTT-AG
yggh_D: CGC-CAT-AAA-GCG-TTC-AAA-AT

k1: CAG-TCA-TAG-CCG-AAT-AGC-CT
k2: CGG-TGC-CCT-GAA-TGA-ACT-GC

rit1_C: TCT-AGG-AAA-AGT-GAG-TTC-AGG-CTT-A
rit1_D: TCG-TGT-AAA-ACC-TCA-GAA-CAC-TGT-A

dus3_C: GGT-CTA-GGC-AAC-AAA-GGT-ACA-CTA-A
dus3_D: TAT-TTT-GAT-TTT-CTT-GGA-ACC-CAT-A

trm10_C: TAC-CAA-TTA-TGA-AAA-CTG-GAA-CCA-T
trm10_D: TAC-AAC-ATC-AAA-GCA-AAA-TAA-GCA-A

kanC: TGA-TTT-TGA-TGA-CGA-GCG-TAA-T

T7 promoter: 5′TAA-TAC-GAC-TCA-CTA-TAG 3′

yeast tRNA$^{Phe}$ DNA template:
3′ ATTATGCTGAGTGATATCGGGCCTATCGAGTCAGCCATCTCGTCCCCTAACTTTTAGGGGCACAGGAACCAAGCTAAGGCTCAGGCCCGTGGT 5′

yeast tRNA$_i^{Met}$ DNA template:
3′ ATTATGCTGAGTGATATCCGCGGCACCGCGTCACCTTCGCGCGTCCCGAGTATTGGGACTACAGGAGCCTAGCTTTGGCTCGCCGCGGTGGT 5′
*Remark: The A1-U72 base pair was replaced with a G1-C72 base pair to improve in vitro transcription efficiency.*

reverse complementary oligo for yeast tRNA$_i^{Met}$ purification:
5′ [Biotin]-AAA-TCG-GTT-TCG-ATC-CGA-GGA-CAT-CAG-GGT-TAT-GA 3′

**Supplementary Table S3**

| Enzyme | Trm2 | | Trm6/Trm61 | | | | | |
|---|---|---|---|---|---|---|---|---|
| tRNA quantity | 500 pmol | | 500 pmol | | | | | |
| Yeast tRNAs | unmodified - $tRNA^{Phe}$ | Ψ55- $tRNA^{Phe}$ | unmodified - $tRNA^{Phe}$ | T54- $tRNA^{Phe}$ | Ψ55- $tRNA^{Phe}$ | Ψ55- T54- $tRNA^{Phe}$ | unmodified- $tRNA_i^{Met}$ | $m^5C48,49$- $tRNA_i^{Met}$ |
| Enzyme (pmol) | 20 | 2.5 | 15 | 15 | 5 | 2.5 | 7.5 | 7.5 |
| SAM (pmol) | 880 | 880 | 880 | 880 | 880 | 880 | 880 | 880 |
| $[^3H]$-SAM (pmol) | 2.5 | 2.5 | 2.5 | 2.5 | 2.5 | 2.5 | 2.5 | 2.5 |
| Reaction conditions | MB1X 30°C 30 min | MB1X 30°C 30 min | MB1X 30°C 96 min | MB1X 30°C 30 min | MB1X 30°C 30 min | MB1X 30°C 30 min | MB1X 30°C 30 min | MB1X 30°C 30 min |

The different kinetic reaction mixes and conditions to determine the initial velocities of Trm2 introducing the T54 modification and Trm6/Trm61 introducing the $m^1A58$ modification to a set of specifically modified yeast tRNAs. MB corresponds the maturation buffer: 100 mM $NaH_2PO_4/K_2HPO_4$ pH 7.0, 5 mM $NH_4Cl$, 2 mM DTT and 0.1 mM EDTA. These quantities correspond to a volume of 50 µL of reaction.

### SUPPLEMENTARY FIGURES

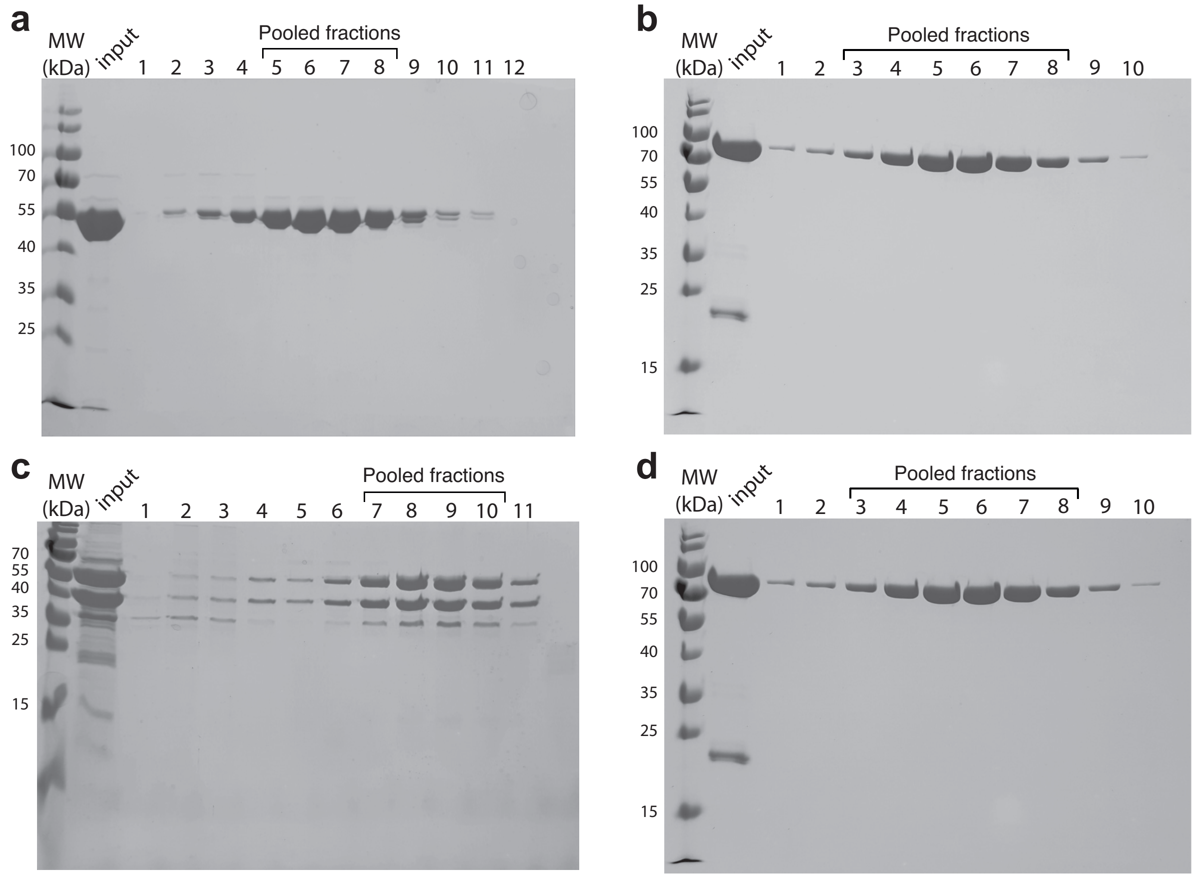


**Supplementary Figure S1: Quality control in the last step of purification of the modification enzymes**

(**a**) SDS-PAGE of the fractions of the Superdex 75 size exclusion column of Pus4 purification. (**b**) SDS-PAGE of the fractions of the Superdex 200 size exclusion column of Trm2 purification. (**c**) SDS-PAGE of the fractions of the Superdex 200 size exclusion column of Trm6/Trm61 purification. (**d**) SDS-PAGE of the fractions of the Superdex 200 size exclusion column of Trm4 purification. (MW): protein ladder, size in kDa. (input): fraction injected onto the column. For each purification, the pooled fractions are indicated above the gel.

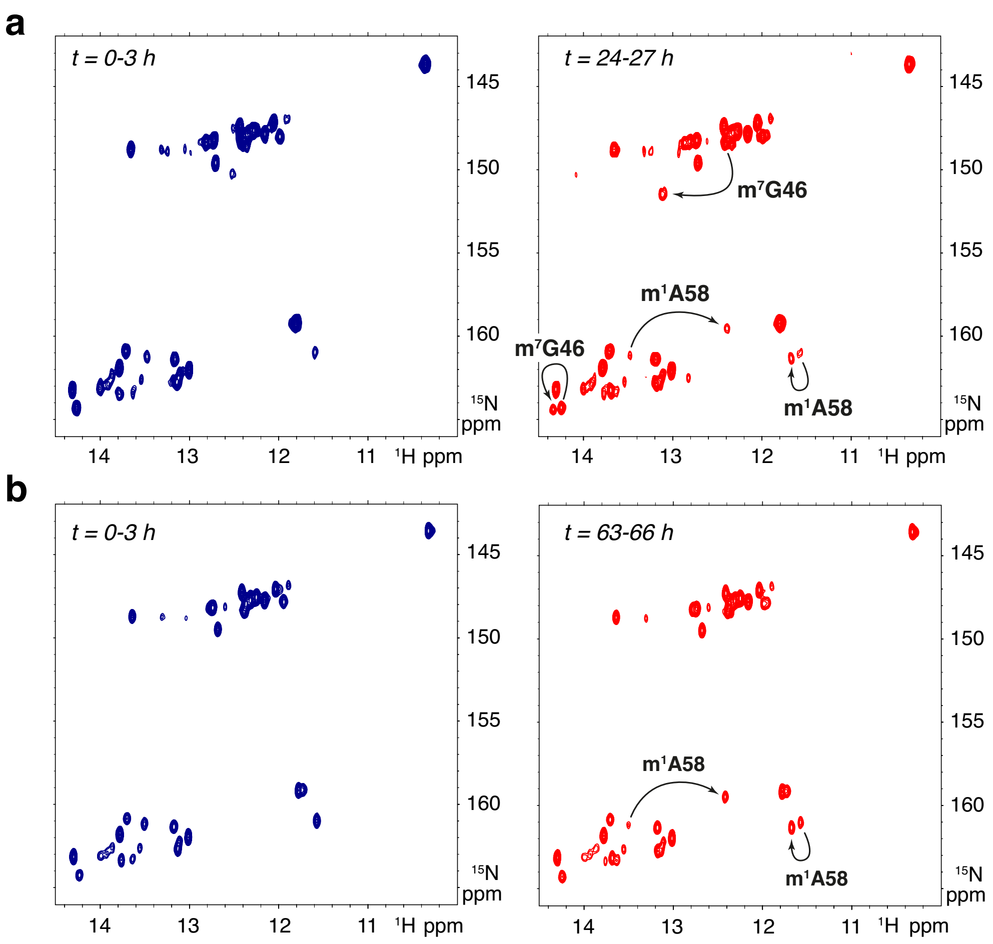


**Supplementary Figure S2: Contamination of Trm6/Trm61 with m$^7$G46 activity**
(**a**) ($^1$H,$^{15}$N)-BEST-TROSY experiments on $^{15}$N-labelled tRNA$^{Phe}$ reporting on Trm6/Trm61 enzymatic activity purified from *E. coli* BL21(DE3)CodonPlus-RIL cells. Concentrations: tRNA$^{Phe}$ at 100 µM and Trm6/Trm61 at 2 µM. Although the m$^1$A58 activity is clearly observable, the Trm6/Trm61 sample is contaminated with m$^7$G46 activity. (**b**) ($^1$H,$^{15}$N)-BEST-TROSY experiments on $^{15}$N-labelled tRNA$^{Phe}$ reporting on Trm6/Trm61 enzymatic activity purified from *E. coli* BL21(DE3)CodonPlus-RIL *yggh::kan* cells. The sample is no longer contaminated with m$^7$G46 activity.

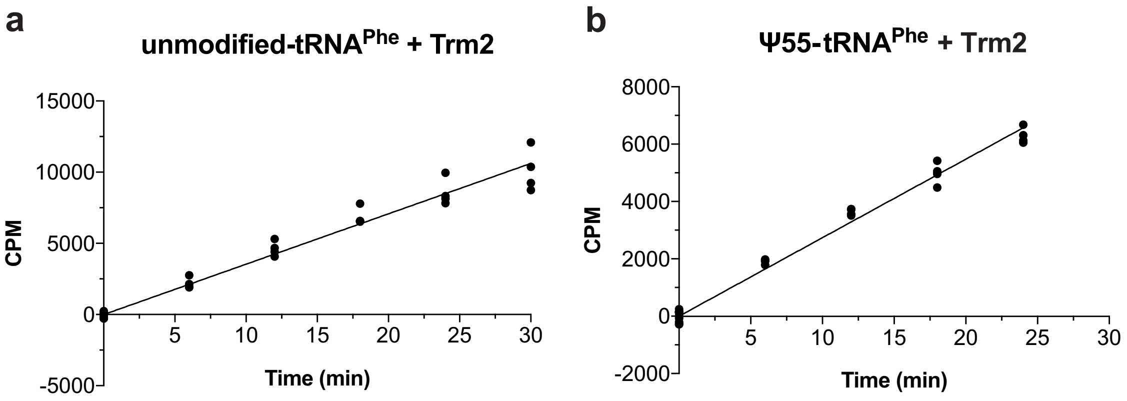


**Supplementary Figure S3: Kinetic efficiency of Trm2 depending on the modification profiles of yeast tRNA$^{Phe}$**

(**a**) Raw counts per minute (CPM) data corresponding to T54 introduction by Trm2 on the yeast unmodified-tRNA$^{Phe}$. (**b**) Raw CPM data corresponding to T54 introduction by Trm2 on the yeast tRNA$^{Phe}$ containing the Ψ55 modification (Ψ55-tRNA$^{Phe}$). CPM are measured for 4 or 5 time points in four independent experiments (N=4). Black dots represent individual measurements.

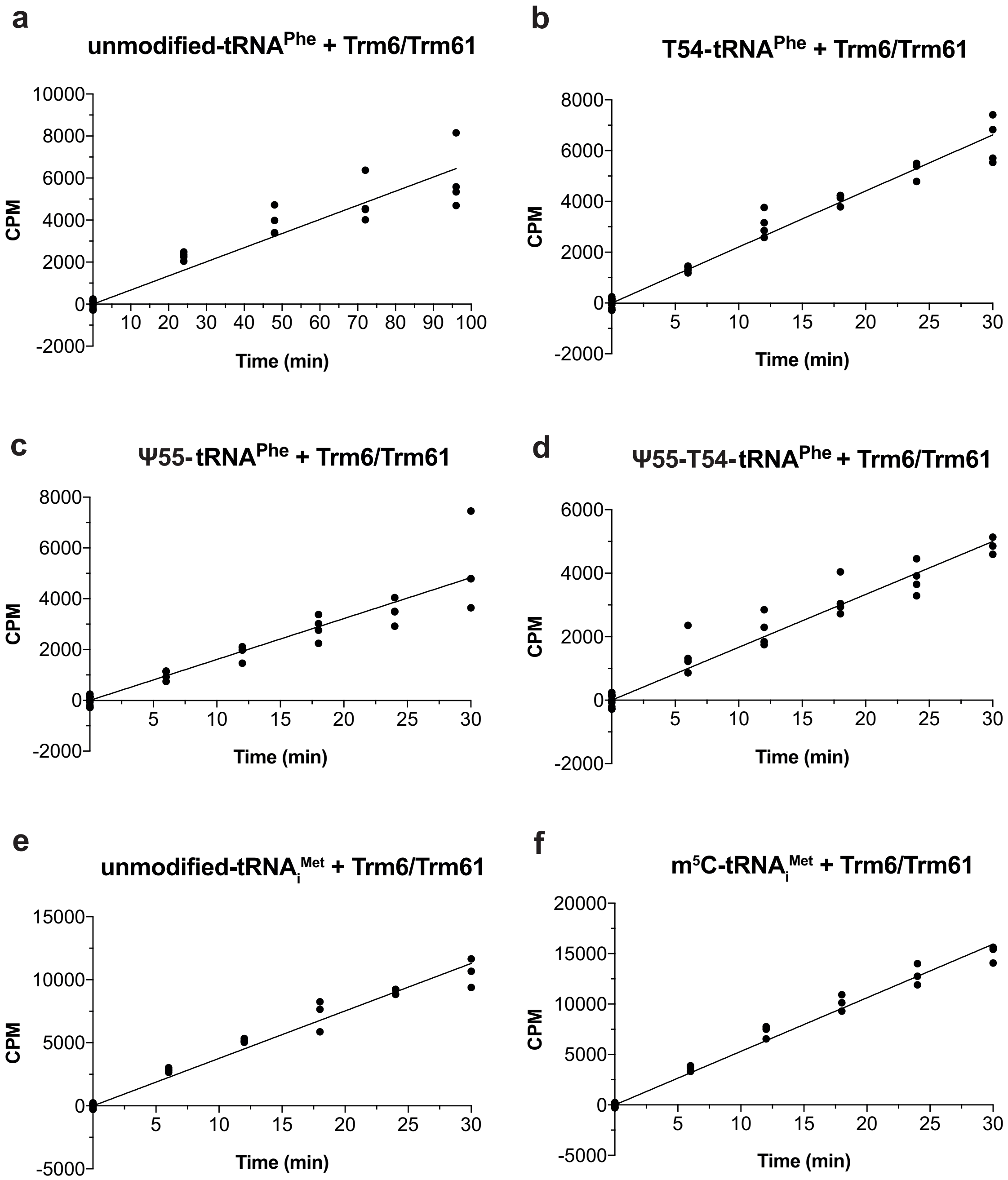


**Supplementary Figure S4: Kinetic efficiency of Trm6/Trm61 depending on the modification profiles of yeast tRNA$^{Phe}$ and tRNA$_i^{Met}$.**

(**a**) Raw counts per minute (CPM) data corresponding to the m$^1$A58 introduction by Trm6/Trm61 on the yeast unmodified-tRNA$^{Phe}$. (**b**) Raw CPM data corresponding to the m$^1$A58 introduction by Trm6/Trm61 on the tRNA$^{Phe}$ containing the T54 modification (T54-tRNA$^{Phe}$). (**c**) Raw CPM data corresponding to the m$^1$A58 introduction by Trm6/Trm61 on the tRNA$^{Phe}$ containing the Ψ55 modification (Ψ55-tRNA$^{Phe}$). (**d**) Raw CPM data corresponding to the m$^1$A58 introduction by Trm6/Trm61 on the tRNA$^{Phe}$ containing both the Ψ55 and T54 modifications (Ψ55-T54-tRNA$^{Phe}$). (**e**) Raw CPM data corresponding to the m$^1$A58 introduction by Trm6/Trm61 on the unmodified-tRNA$_i^{Met}$. (**f**) Raw CPM data corresponding to the m$^1$A58 introduction by Trm6/Trm61 on the tRNA$_i^{Met}$ containing the m$^5$C48,49 modifications (m$^5$C-tRNA$_i^{Met}$). CPM are measured for 4 or 5 time points in at least three independent experiments (N=3 or 4). Black dots represent individual measurements.
